# Atomic structures of medin and Aβ fibrils reveal polymorphic remodeling in mixed amyloid systems

**DOI:** 10.1101/2025.07.17.665283

**Authors:** Brajabandhu Pradhan, Senthil T Kumar, Jessica Wagner, Rodrigo Gallardo, Gabriele Orlando, Matthias De Vleeschouwer, Valentina zorzini, Jillian Madine, Nikolas Louros, Jonas J. Neher, Frederic Rousseau, Joost Schymkowitz

**Affiliations:** Switch Laboratory, VIB Center for Brain and Disease Research, 3000 Leuven, Belgium; Switch Laboratory, Department of Cellular and Molecular Medicine, KU Leuven, 3000 Leuven, Belgium; Biomedical Center (BMC), Biochemistry, Faculty of Medicine, LMU Munich, Munich, Germany; Neuroimmunology and Neurodegenerative Diseases, German Center for Neurodegenerative Diseases (DZNE), Munich, Germany; Department of Cellular Neurology, Hertie Institute for Clinical Brain Research, University of Tübingen, Tübingen, Germany; Munich Cluster for Systems Neurology (SyNergy), Munich, Germany; Institute of Systems, Molecular & Integrative Biology, University of Liverpool, United Kingdom

## Abstract

Amyloid assembly in vivo occurs in complex proteomic environments where multiple aggregation-prone species coexist. Aβ and medin are two highly prevalent amyloids in ageing humans, and medin co-localizes with Aβ in cerebral amyloid angiopathy (CAA). Yet the structural consequences of their interaction remain unclear. Here, we determine the cryo-electron microscopy (cryo-EM) structures of fibrils formed from defined in vitro mixtures of Aβ40 and medin. From the same reaction, we resolve three fibril populations: (i) a previously described Aβ40 polymorph that also forms in isolation, (ii) a novel Aβ40 polymorph with Aβ42-like features, including ordered N- and C-terminal regions, and (iii) the first atomic structure of full-length medin fibrils. Biochemical and immunogold analyses demonstrate association between Aβ and medin within the mixed assemblies. However, the cryo-EM map of the novel Aβ polymorph does not unambiguously resolve medin within the ordered Aβ core. These findings are consistent with two non-exclusive mechanisms by which the polymorphic landscape is reshaped in mixtures: transient heterotypic amyloid interactions that redirect Aβ folding pathways, or partial and structurally heterogeneous incorporation of medin into the fibril architecture. Regardless of the precise structural mechanism, our data demonstrate that interactions between coexisting amyloids can remodel their polymorphic landscape, revealing how mixed amyloid systems generate emergent conformations.

## Introduction

Amyloid-β (Aβ) fibrils are key pathological drivers in Alzheimer’s disease (AD) and cerebral amyloid angiopathy (CAA), where they accumulate in parenchymal plaques and cerebral vessel walls, respectively (Selkoe and Hardy 2016). Aβ peptides - particularly Aβ40 and Aβ42 - self-assemble into structurally polymorphic fibrils, and their conformations are shaped by the biochemical environment in which they form (Tycko 2015, Gallardo, Ranson et al. 2020). These polymorphic states are thought to influence properties such as seeding behavior, tissue distribution, cell toxicity, and disease progression (Tycko 2015, Gallardo, Ranson et al. 2020, Li and Liu 2023). However, the molecular determinants governing this structural diversity remain incompletely understood.

One emerging principle is that heterotypic interactions - where Aβ co-aggregates with other amyloidogenic peptides or cofactors - can modulate fibril morphology, aggregation kinetics, and strain identity (Konstantoulea, Guerreiro et al. 2022, Louros, Ramakers et al. 2022, Louros, Schymkowitz et al. 2022). Recent cryo-EM studies have demonstrated that distinct amyloidogenic proteins can co-assemble into heteromeric filaments in vivo, most notably in the case of TDP-43 and Annexin-11 filaments extracted directly from patient tissue (Arseni, Nonaka et al. 2024). These findings establish that heterotypic amyloid interactions can yield fully ordered mixed fibril cores. However, whether such interactions always result in stable heteromeric architectures or can instead act transiently to reshape the polymorphic landscape of one partner without permanent incorporation into the ordered core remains unclear.

A prominent example is medin, a 50-residue cleavage product of milk fat globule-EGF factor 8 (MFG-E8), and the most prevalent localized human amyloid (Mucchiano, Cornwell et al. 1992, Madine, Davies et al. 2023). Medin deposits in the vasculature of nearly all individuals over 50 years and contributes to vascular stiffening and dysfunction (Häggqvist, Näslund et al. 1999, Degenhardt, Wagner et al. 2020). It is a defining feature of age-related aortic medial amyloidosis and we previously showed that it co-localizes and co-aggregates with Aβ in both AD patients (with CAA) and transgenic mouse model (Degenhardt, Wagner et al. 2020, Wagner, Degenhardt et al. 2022). In particular, in APP23 and APP Dutch mice, which develop substantial vascular Aβ deposition (Sturchler-Pierrat, Abramowski et al. 1997, Herzig, Winkler et al. 2004), we found that medin co-deposits with Aβ (Wagner, Degenhardt et al. 2022). Moreover, in APP23 mice medin exacerbated cerebrovascular pathology as genetic ablation of the medin-containing C2 domain of its precursor protein MFG-E8 substantially reduced vascular Aβ burden. Notably, we found that the presence of medin may alter Aβ fibril structure, as indicated by amyloid conformation-sensitive dyes (Wagner, Degenhardt et al. 2022). However, the exact structural changes resulting from this amyloid-amyloid interaction remain unknown.

With the growing interest in medin and its interplay with Aβ in ageing-associated vascular amyloidosis and pathology (Madine, Davies et al. 2023), understanding the structural basis of medin fibril formation is essential. However, no high-resolution structure of full-length medin fibrils has been reported to date. Solid-state NMR analysis of the 50-residue peptide has indicated the presence of extended β-strands and a salt bridge between Asp25 and Lys30 (Davies, Madine et al. 2015). Similarly, solid-state NMR and X-ray diffraction studies of C-terminal fragments comprising the aggregation-prone region (residues 42–49) have shown that these segments form amyloid-like aggregates (Madine, Copland et al. 2009).

However, these findings do not provide a high-resolution description of the molecular architecture of full-length medin fibrils, nor do they account for the morphological heterogeneity observed in assembled fibrils (Larsson, Söderberg et al. 2007). A comprehensive structural framework for full-length medin, therefore, remains lacking.

Here, we investigate the structural consequences of heterotypic co-aggregation between recombinantly purified Aβ40 and synthetic medin peptides in vitro. Using cryo-EM, aggregation kinetics, biophysical assays, and immunogold electron microscopy, we find that medin accelerates Aβ40 aggregation and promotes the emergence of distinct fibril polymorphs. Cryo-EM analysis reveals two coexisting fibril polymorphs of Aβ40: one matching a previously described in vitro Aβ40 structure, and a second, previously unreported morphology with structured N- and C-terminal regions and Aβ42-like architectural features. In addition, we resolve the first atomic structure of full-length medin fibrils from the same mixed assembly reaction. The novel Aβ40 polymorph reproducibly emerges in mixtures but not in isolated Aβ40 samples under comparable conditions. Although biochemical and immunogold analyses demonstrate an association between Aβ and medin within the mixed assemblies, the cryo-EM map does not unambiguously resolve medin within the ordered Aβ core. These findings are therefore consistent with a spectrum of heterotypic outcomes: from stable co-assembly within a shared fibril architecture, as observed in TDP-43/Annexin-11 filaments, to transient or structurally heterogeneous interactions that redirect the folding pathway of one amyloidogenic partner without fully ordered incorporation.

Although medin and Aβ are known to co-deposit in vivo - particularly in animal models and human cases characterized by robust vascular Aβ deposition (Wagner, Degenhardt et al. 2022) - the fibril cryo-EM structures reported here differ from known ex vivo CAA fibrils (Kollmer, Close et al. 2019, Yang, Murzin et al. 2023). In vivo, medin and Aβ may aggregate at different times and encounter each other as partially assembled or mature fibrils.

However, isolating fibrils of co-aggregated Aβ and Medin from human samples for structural determination remains technically challenging. Therefore, the current study examines structural outcomes arising from full co-aggregation of Aβ40 and medin monomers in vitro. Our findings suggest that heterotypic amyloid interactions operate along a continuum of structural possibilities and can reshape the conformational landscape of Aβ fibrils even when the interacting partner is not fully resolved within the final ordered core.

Moreover, we report the first cryo-EM structure of full-length medin fibrils, identified in one of our co-aggregated Aβ–medin samples. The structure reveals an ‘r’ shaped nonplanar fold of medin in which the polar residue-rich N-terminus forms the fibril core, while the C-terminal aggregation-prone region is exposed to the solvent. Notably, the structure contains a rare polar steric zipper formed between residues from the N- and C-terminal halves of adjacent subunits, distinguishing medin from many previously characterized amyloids that are predominantly stabilized by hydrophobic interfaces. This first atomic model of full-length medin provides a molecular framework for understanding its aggregation behavior.

## Results

### Heterotypic co-aggregation with medin alters Aβ40 aggregation kinetics

Thioflavin T (ThT) fluorescence assays were performed to monitor the co-aggregation kinetics of synthetic Aβ40 and medin. Monomeric Aβ40 (20 µM), medin (20 µM), and a mixture of Aβ40 (20 µM) and medin (20 µM) were dissolved in a buffer containing 50 mM Tris and 100 mM NaCl, pH 7.4. The samples were incubated at 30°C, and shaken every 10 minutes, with fluorescence recorded over a period of five days.

Under these experimental conditions, we observed that Aβ40 has a longer lag phase of 23 hours compared to medin, which has a lag phase of only approximately 3 hours (Fig. 1A). However, in the mixture of Aβ40 and medin, the lag phase is 6.5 hours, suggesting that the presence of medin accelerates the aggregation of Aβ40 (Fig. 1A). These observations are consistent with our previous study demonstrating that medin directly promotes Aβ40 aggregation (Wagner, Degenhardt et al. 2022). Although the lag times observed here are longer than those reported previously, this difference is attributable to the lower incubation temperature (30 °C) and the use of a different buffer composition compared to our earlier study (150 mM NaCl, 5 mM EDTA). Representative kinetic traces from six independent experiments are shown in Supplementary Fig. 1.

**Figure 1:**
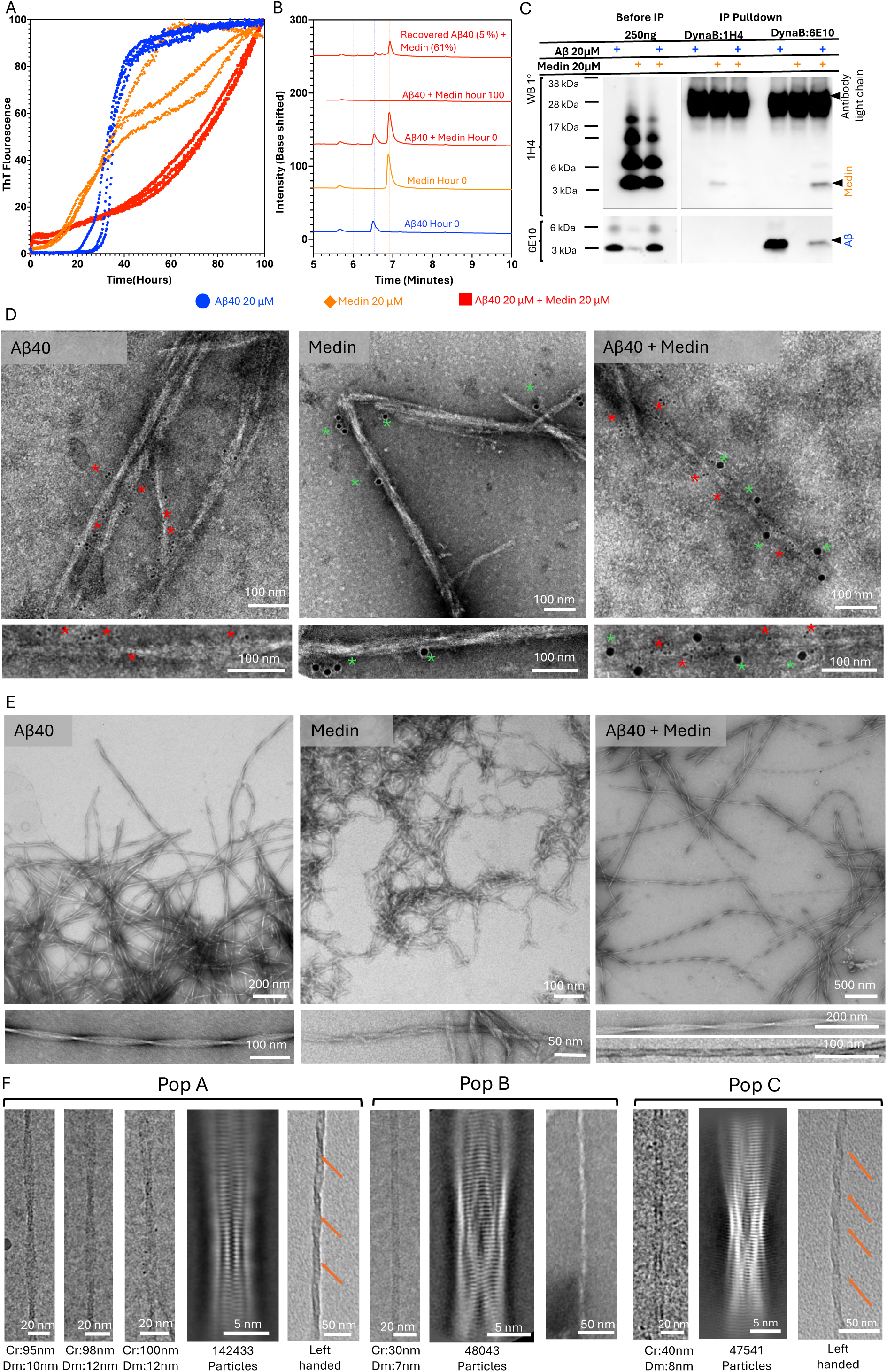
Medin accelerates Aβ40 aggregation and remodels fibril morphology through heterotypic co-assembly. **(A)** Thioflavin T (ThT) fluorescence kinetics of Aβ40 (20 µM), medin (20 µM) and an equimolar mixture in 50 mM Tris, 100 mM NaCl, pH 7.4 at 30 °C. Aβ40 exhibits a lag phase of ∼23 h, whereas medin aggregates within ∼3 h. The mixture shows an intermediate lag time (∼6.5 h), indicating accelerated Aβ40 nucleation in the presence of medin. Data represent three technical replicates (n = 3), normalized to the maximal fluorescence of each replicate. **(B)** LC–MS chromatograms of monomeric (0 h) and post-aggregation (5 d) samples. Peaks corresponding to Aβ40 (6.5 min) and medin (6.9 min) are absent after aggregation and reappear following HFIP solubilization, confirming incorporation of both peptides into fibrils. Chromatograms (except Aβ40 at 0 h) were baseline-shifted for clarity. **(C)** Co-immunoprecipitation of fibrils formed from Aβ40, medin, or co-aggregated Aβ40– medin using anti-Aβ (6E10) or anti-medin (1H4) antibodies coupled to Dynabeads, followed by immunoblotting with 1H4 (top) and 6E10 (bottom). Pull-down with 6E10 recovers both Aβ and medin from co-aggregated samples, demonstrating heterotypic assembly. Reduced Aβ signal compared to Aβ-only fibrils suggests partial masking of the N-terminal 6E10 epitope. Immunoprecipitation with 1H4 is inefficient, consistent with lower apparent affinity. **(D)** Dual immunogold labelling of fibrils with anti-Aβ (6E10; 6 nm gold) and anti-medin (1H4; 10 nm gold). Aβ40 and medin fibrils alone show exclusive labelling, whereas mixed samples display singly and doubly labelled fibrils, indicating coexisting homotypic (Supplementary Fig. 2) and heterotypic assemblies. **(E)** Negative-stain TEM images show that Aβ40 forms heterogeneous ribbon-like fibrils and medin forms morphologically diverse filaments lacking defined crossover. In contrast, Aβ40–medin fibrils exhibit pronounced helical twist and reduced heterogeneity. **(F)** Representative cryo-EM micrograph and corresponding 2D class averages of Aβ40–medin fibrils reveal three distinct populations: population A, wide twisted fibrils with ∼100 nm crossover; population B, narrow straight fibrils with ∼30 nm crossover (experiment 1); and population C, fibrils with ∼40 nm crossover and ∼90 Å diameter (experiment 3). Chromium shadowing (shown to the right of the 2D classes) indicates that populations A and C are left-handed, whereas the handedness of population B could not be unambiguously determined owing to the absence of clearly resolved crossovers. Handedness is indicated by orange arrows.

To further validate the co-aggregation of Aβ40 and medin, LC-MS analysis was performed on the peptides before and after the ThT assay. The concentration of unused monomeric peptides (Aβ40 and medin) was monitored at various time points during the 5-day incubation period to track their consumption and the formation of fibrils (Fig. 1B, Supplementary Fig. 2). Prior to aggregation, we observed LC peaks at 6.5 minutes corresponding to Aβ40 and at 6.9 minutes corresponding to medin (initiation phase) (Fig. 1B). After 5 days, 100% of the monomers were consumed (saturation phase), as indicated by the lack of medin or Aβ40 peaks (Fig. 1B). Interestingly, quantifying the monomer concentration over the course of the ThT assay revealed that monomers remained detectable in both the Aβ40-only and medin-only untill day two and three respectively (Supplementary Fig. 2A). In contrast, the Aβ40+medin sample exhibited a pronounced depletion of monomers after day one, indicating an accelerated monomer consumption in the co-aggregated sample (Supplementary Fig. 2A). A sample from the saturation phase was centrifuged at 21000x g to recover the fibrils and was dissolved with 95% HFIP. Subsequent LC-MS analysis of the dissolved material showed the reappearance of peaks corresponding to both Aβ40 and medin (Fig. 1B). Notably, HFIP dissolution allowed the recovery of approximately 61% of medin monomers from the co-aggregated fibrils, a significantly higher proportion than that of Aβ40 (5 %) (Fig. 1B), suggesting greater resistance of pure Aβ40 fibrils to HFIP mediated disassembly.

To investigate the heterotypic nature of the aggregates, we performed co-immunoprecipitation experiments. Fibrils were separated by centrifugation at 21,000 rcf. Samples containing Aβ40 fibrils, medin fibrils, and Aβ40-medin co-aggregated fibrils were subjected to pull-down using either the anti-medin antibody 1H4, a monoclonal antibody targeting the N-terminus of medin (Degenhardt, Wagner et al. 2020). or the anti-Aβ antibody 6E10 which binds to residues 3-8 (EFRHDS) of Aβ, both coupled to magnetic Dynabeads. After incubation, the bound fibrils were denatured, digested, and sonicated, followed by Western blot analysis. The 6E10 antibody successfully pulled down both Aβ and medin in the Aβ–medin co-aggregated sample, confirming their heterotypic association (Fig. 1C).

However, the intensity of the Aβ band in the co-aggregated sample was significantly lower compared to the Aβ-only sample. This reduced binding can be explained by partial masking of the 6E10 epitope, which lies in the N-terminus of Aβ40, a region likely involved in interaction with medin. As a result, 6E10 binds only where the epitope remains exposed. In contrast, pull-down using the medin antibody 1H4 was not successful. This is most likely due to the relatively lower affinity of the antibody, as indicated by the faint medin band observed even in the medin-only sample.

To determine whether Aβ40 and medin co-assemble into heterotypic fibrils, we performed dual immunogold electron microscopy (Immunogold EM) using primary antibodies 6E10, (Aβ40: 6nm gold particles), and 1H4 (medin:10 nm gold particles), Immunogold EM images of Aβ40 and medin fibrils alone showed no cross-reactivity of the antibodies, confirming their specificity (Fig. 1D). Notably, Aβ40 fibrils exhibited a significantly higher frequency of 6 nm gold particles (6E10) compared to the 10 nm gold particles (1H4) labeling medin fibrils, indicating greater binding efficiency of 6E10 than 1H4 (Fig. 1D). In fibrils formed from the Aβ40–medin mixture, we observed fibrils labeled with 6E10 alone (indicative of homotypic Aβ40 fibrils: Supplementary Fig. 3A), 1H4 alone (indicative of homotypic medin fibrils: (Supplementary Fig. 3B), and both 6E10 and 1H4 (Fig. 1D), demonstrating the formation of heterotypic Aβ40:medin fibrils.

### Medin Reshapes the Polymorphic Landscape of Aβ40 Fibrils

The fibril morphology of saturation-phase samples was first examined using negative-stain TEM (NS-TEM) and cryo-EM. Medin alone self-assembled into a highly heterogeneous mixture of fibrils, including straight, rigid filaments and flexible, noodle-like structures. These fibrils lacked a well-defined helical twist and displayed variable diameters (Fig. 1E). In contrast, Aβ40 alone formed heterogeneous, straight ribbon-like filaments that were occasionally twisted and frequently laterally associated (Fig. 1E). Strikingly, fibrils formed by co-aggregation of Aβ40 and medin exhibited well-defined helical crossovers and markedly reduced morphological heterogeneity (Fig. 1E).

To quantify fibril polymorphism, cryo-EM analysis was performed on Aβ40 alone (two independent datasets) and Aβ40–medin co-aggregation samples (three independent datasets). Morphology composition was quantified by counting particles in cryoSPARC 2D class averages.

In the first Aβ40-alone dataset (1,766 micrographs), most fibrils were laterally bundled and unsuitable for cryo-EM helical reconstruction. Only ∼5% displayed discernible helical twisting (Morphologies I and II; Supplementary Fig. 4A). By contrast, the corresponding Aβ40–medin dataset (7,302 micrographs) showed a substantial enrichment of well-ordered, twisted fibrils. Approximately 85% of particles exhibited clear helical crossovers, while ∼15% were bundled and excluded from further analysis. Three morphologies were identified based on 2D class averages. Morphology I (58.79%) and Morphology II consisted of well-twisted fibrils with variable diameters of 10–12 nm and crossover distances of 90–100 nm (Fig. 1F; Supplementary Fig. 4B). Morphology III (9.44%) comprised straight fibrils with a uniform diameter of ∼6 nm and a fixed crossover distance of ∼30 nm. High-resolution reconstructions were obtained for Morphologies I and III from this dataset, hereafter termed Population A and Population B, respectively. Population B represents a novel Aβ40 fibril morphology with a distinct unidentified cryo-EM density, as described in the following sections.

A second independent Aβ40-medin aggregation experiment confirmed the reproducibility of these polymorphs. Populations A and B were again observed (Morphologies I and III; Supplementary Fig. 4C), each exhibiting helical parameters consistent with those from the first experiment. However, the proportion of particles corresponding to Population B decreased to 5.79%. Interestingly, an additional twisted morphology was detected, characterized by a crossover distance of ∼50 nm and a diameter of ∼8 nm, representing approximately 6.16% of particles. Owing to the lower overall data quality and the limited number of particles for these minor morphologies, high-resolution structures could not be determined.

To enhance the representation of rare polymorphs, a third experiment was conducted using 2-nm-thick continuous carbon support films instead of the holey carbon grids employed in the previous experiments. Under these conditions, the Aβ40-alone dataset (7,056 micrographs) showed a marked increase in twisted fibrils (∼44%), of which 12.81% resembled Population A identified in the first experiment (Supplementary Fig. 4E). Notably, approximately 31% of the twisted fibrils exhibited larger diameters (>16 nm) and extended crossover distances (∼116 nm). The remaining fibrils were highly heterogeneous and could not be reliably classified into distinct morphologies.

The corresponding Aβ40-medin mixture dataset (15,891 micrographs) displayed an expanded polymorphic landscape, including twisted fibrils with increased diameters and extended crossover distances. Nearly 18% of fibrils exhibited diameters ≥15 nm and crossover distances >115 nm, comparable to those observed in the Aβ40-alone sample. Population A fibrils were observed in both samples in this experiment, albeit at a lower proportion (4.2%) in the Aβ40-medin mixture.

In addition, ∼18% of particles (Morphology I in Supplementary Fig. 4F) corresponded to a distinct fibril morphology not observed in previous datasets. These fibrils are characterized by a diameter of ∼8 nm and a crossover distance of ∼40 nm. High-resolution cryo-EM analysis revealed that this morphology consisted exclusively of medin. This structure is hereafter referred to as Population C (Fig. 1F).

### Population A: Aβ40 fibrils with a four-layered cross-β structure

Helical reconstruction of Population A fibrils yielded a three-dimensional (3D) cryo-EM map at 3.1 Å resolution, based on the Fourier shell correlation (FSC) 0.143 criterion (Supplementary Fig. 5A). The fibrils display well-resolved cross-β layers with clear side-chain density and consist of two protofilaments related by pseudo-2₁ screw symmetry (Fig. 2A). Each cross-section exhibits a characteristic four-layered architecture, comprising two extended inner β-layers and two shorter, outer β-layers (Figs. 2A, B). Chromium side-shadowing experiments revealed a left-handed twist in these fibrils (Fig. 1F).

**Figure 2:**
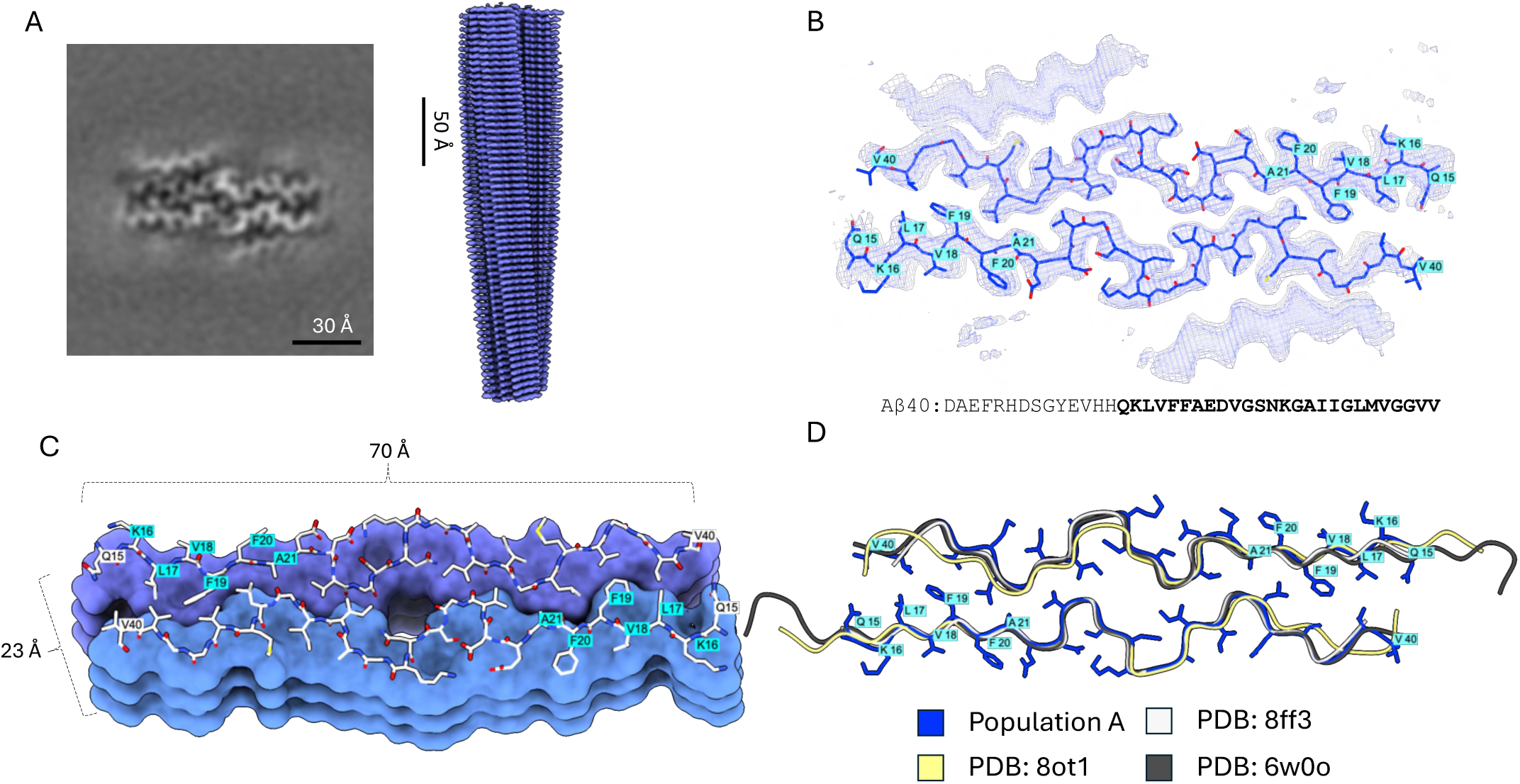
Cryo-EM structure of Population A fibrils reveals a four-layered cross-β architecture. **(A)** Cryo-EM density map of Population A fibrils (right) and transverse section (left) revealing a four-layered cross-β arrangement. **(B)** Transverse section of the 3D cryo-EM map showing two inner β-sheets composed of extended β-strands spanning residues Q15–V40. The structure was reconstructed from a single cryo-EM dataset derived from one replicate of the ThT assay (n = 1). Map statistics: mean 2.902 × 10⁻⁵, s.d. (σ) 0.001057, and RMS 0.001058. The map is visualized at a contour level of 7σ. **(C)** Surface representation of the atomic model of Population A, highlighting residues within the aggregation-prone region and the interface between the two protofibrils. **(D)** Structural comparison of Population A with previously reported Aβ40 fibril structures (PDB IDs: 8FF3, 8OT1, and 6W0O). Top, cross-sectional alignment; bottom, side-view alignment. In the Population A model, side chains are shown as sticks, whereas reference fibrils are displayed as backbone traces only, each coloured individually as indicated.

An atomic model was built into the inner layers of the map, which provided sufficient side-chain resolution (Figs. 2B, C). The core comprises residues 15-40 of Aβ40, arranged into three β-strands: a longer strand (H14-E22) encompassing the aggregation-prone region KLVFFA, and two shorter strands (A30-I32 and L34-V36) separated by glycine residues G33 and G37 (Fig. 2B). A structured loop formed by residues 21-28 (DVGSNK) is stabilized by a salt bridge between D23 and K28. The protofilament interface is stabilized primarily through inter-subunit hydrophobic interactions, while no clear intra-subunit hydrophobic packing was observed. Density corresponding to the N-terminal region (residues 1-14) was not resolved, likely due to conformational flexibility. The outer β-layers, while visible as peptide-like density, lacked resolvable side chains and thus were excluded from model building.

This Aβ40 conformation has been previously reported in multiple cryo-EM studies of seeded and unseeded fibril growth. Ghosh et al. (2021) observed this fold in Aβ40 in vitro fibrils seeded with human parenchymal AD plaques (pdb : 6w0o) (Ghosh, Thurber et al. 2021). Fu et al. (2024) resolved an identical structure from in vitro Aβ40 fibrils seeded with vascular Aβ deposits derived from Dutch-type CAA patient brains (pdb : 8ff3) (Fu, Crooks et al. 2024) and Pfeiffer et al. (2024) described a similar morphology with minor variations in the C-terminal residues V37–V39 (pdb : 8ot4) (Pfeiffer, Ugrina et al. 2024) again in Aβ40 in vitro fibrils seeded with CAA patient material. Structural alignment revealed strong conservation of the inner β-sheet backbone across all cases (Fig. 2D). The two outer densities visible in our reconstruction also appear in those studies (Supplementary Fig. 6) and were previously attributed to β-hairpins formed by Aβ itself (Ghosh, Thurber et al. 2021), rather than heterotypic sequences such as medin.

Taken together, these findings suggest that Population A represents a conserved and recurrent Aβ40 polymorph that frequently emerges under in vitro conditions, including seeded and unseeded fibril growth. Notably, the same structure is observed here in the presence of medin and has also been reported in multiple Aβ40 seeding experiments using patient-derived parenchymal and vascular amyloid material. This strongly suggests that the fold does not reflect the structure of the original in vivo seeds, but instead represents a common structural endpoint formed de novo during in vitro elongation. Indeed, Pfeiffer et al. (2024) demonstrated that the fibril structure formed during in vitro seeding with CAA-derived material (pdb : 8ot4) differs from the structure of the original seeds themselves (Pfeiffer, Ugrina et al. 2024). Consistent with this and given the absence of sequence-specific evidence for medin in the resolved regions of Population A, we conclude that medin is unlikely to contribute directly to the assembly of this fibril morphology under the conditions tested.

### Population B: Aβ40 Forms a Novel Fibril Polymorph with Structured N- and C-Termini

Helical reconstruction of Population B fibrils using CryoSPARC yielded a 3D cryo-EM map at 2.6 Å resolution, as determined by the FSC 0.143 criterion (Supplementary Fig. 5B). Chromium shadowing experiments to determine the handedness of these fibrils did not yield an unambiguous result. The fibrils adopt a distinctive cuboidal morphology, due to which the helical grooves are not prominently visible in low-resolution TEM images, precluding clear identification of handedness. These fibrils are composed of two twisted protofilaments arranged in a parallel, in-register cross-β architecture (Fig. 3A). The core density is well resolved, allowing for reliable atomic model building and side-chain assignment, and is identified as Aβ(1-40) (Fig. 3B). Surrounding the core, two peripheral faces display low-resolution, fragmented densities resembling polypeptide chains (Fig. 3B). These extra densities do not conform to the fibril symmetry and may correspond to either Aβ segments or co-aggregated medin. Notably, this fibril morphology has not been previously reported for Aβ40.

**Figure 3:**
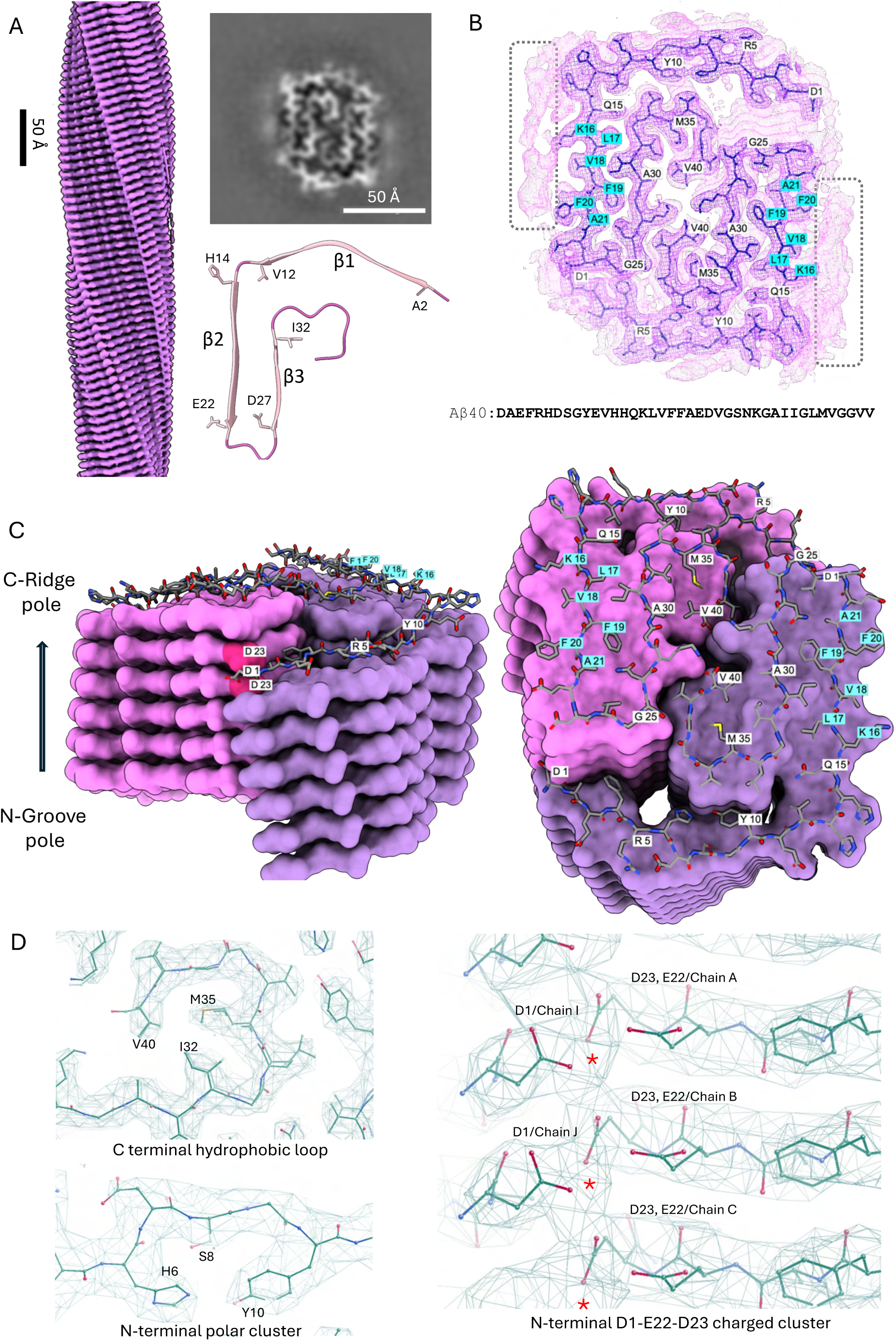
Cryo-EM structure of Population B fibrils reveals a novel F-shaped Aβ40 subunit. **(A)** Cryo-EM density map of Population B fibrils (left) and transverse section highlighting two F-shaped subunits (top right). Cartoon representation of a single F-shaped subunit composed of three β-strands (β1–β3) is shown (bottom right). The structure was reconstructed from a single cryo-EM dataset derived from one replicate of the ThT assay (n = 1). **(B)** Transverse section of the 3D cryo-EM map fitted with two F-shaped subunits comprising Aβ40 (D1–V40). The subunits face each other and form the protofibril interface through C-terminal interactions. Unresolved peripheral densities are indicated by a red box. The map has a mean of 0.0002155, s.d. (σ) of 0.01237 and RMS of 0.01237, with additional densities visible at 3σ. The same map visualized at 7σ is shown in violet. **(C)** Surface representation of the atomic model illustrating the staggered polar arrangement of the two protofibrils. Key residues are annotated, including the amyloidogenic KLVFFA segment. **(D)** Structural features stabilizing Population B fibrils. (Top left) Cryo-EM density of C-terminal residues 28–40 (GAIIGLMVGGVV), forming a hydrophobic loop with V40, I32 and M35 buried in the fibril core. (Bottom left) Density corresponding to the N-terminal polar cluster formed by H6, S8, and Y10. (Right) Interaction of D1 from one protofibril with E22 and D23 of a neighboring subunit from the opposing protofibril, forming a charge cluster. A putative metal ion site is indicated by a red asterisk.

Each protofilament is composed of non-planar, F-shaped subunits that contain three β-strands (Fig. 3A). Strand β1 (A2–V12) and β2 (H14–E22) form extended β-sheets at the periphery of the fibril, while Strand β3 (D27–I32) packs internally and folds back onto Strand 2 (Fig. 3A). Strand 1 forms a tilted β-sheet (∼10° relative to the fibril cross-section), contributing to the cuboidal profile. Due to a ±2 Å axial offset between subunits, the fibril is polar, with one end characterized by a solvent-exposed N-terminal groove and the other by a hydrophobic C-terminal ridge (Fig. 3C).

Stabilization of this structure arises from both hydrophobic and electrostatic interactions. Charged residues in the N-terminal segment (E3, R5, D7, E11) are solvent-exposed, while hydrophobic residues contribute to the internal core. Polar contacts involving H6, S8, and Y10 form a stabilizing cluster near the fibril surface (Fig. 3D). The aggregation-prone KLVFFA motif (residues 16–21) in β strand 2 is partially solvent-exposed, and the adjacent cryo-EM density of unknown identity aligns with residues H13–F19 (Fig. 3B), suggesting a potential interaction site for medin or disordered Aβ segments.

The two protofilaments are joined through a staggered interface dominated by hydrophobic packing and electrostatic contacts. Residues 28–40 (GAIIGLMVGGVV) form a compact, C-shaped hydrophobic loop formed by sharp turns at G33, G37, with I32, M35, and V40 buried in the fibril interior (Fig. 3D). The inter-protofilament interface is further stabilized by hydrogen bonding between K28 (from one protofilament) and the backbone carbonyl of V40 (from the opposing protofilament). Unlike in canonical Aβ40 fibrils, where D23 and E22 form salt bridges, in this morphology, they are solvent-exposed and participate in a distinctive zipper-like charge cluster with D1 from the N-termini of opposing subunits (Fig. 3D). This electrostatic motif may involve additional stabilization by metal ions, although the resolution does not allow unambiguous identification of coordinating species (Fig. 3D).

Taken together, Population B represents a previously unobserved Aβ40 polymorph characterized by ordered N- and C-terminal regions, polar subunit geometry, and a structurally integrated hydrophobic interface. The presence of adjacent unknown densities and the absence of this morphology in Aβ40-alone samples suggest that its formation may be promoted or stabilized by heterotypic interaction with medin.

### Model-Fitting and Energetic Analysis Implicate Aβ in Population A and medin in Population B

To determine the identity of the unassigned cryo-EM densities observed in both fibril morphologies, we employed a combined strategy of model fitting and energy profiling. Although these densities lacked sufficient resolution for de novo atomic modeling, they allowed manual placement of a poly-alanine backbone: 10 residues in Population A and 9 in Population B in Coot (Emsley, Lohkamp et al. 2010). Using FoldX (Schymkowitz, Borg et al. 2005, Delgado, Reche et al. 2025), we generated structural models for all possible peptide segments by applying a sliding window across the sequences of both Aβ40 and medin. For each candidate, we calculated interaction energies and assessed map-to-model agreement with pyUUL (Orlando, Raimondi et al. 2022), quantified as a Pearson correlation coefficient.

For Population A, the resolution of the additional density was sufficient to generate confident predictions. The top-scoring segment was AEDVGSNKGA (residues 22–32 of Aβ), which yielded the most favorable combination of FoldX interaction energy and map correlation (Supplementary Fig. 7). Additional high-scoring sequences - GSNKGAIIGL and GAIIGLMVG - also originated from Aβ40, supporting the conclusion that the density corresponds to an Aβ-derived segment (Supplementary Fig. 7). These results are consistent with prior reports attributing similar densities to β-hairpin structures formed by Aβ40 itself, as observed by (Ghosh, Thurber et al. 2021) and (Pfeiffer, Ugrina et al. 2024) in the context of extended four-layered morphologies containing only Aβ.

We then turned our attention to population B fibrils since our immunogold labeling data had shown co-localization of medin and Aβ40 within individual fibrils (Fig. 1D). The additional density in Population B appeared to be anchored on residues H13, H14, and K16 of Aβ, i.e., close to the crucial central aggregation-prone region. Contacts in this region are likely to contribute to the open N-terminal conformation unique to the F morphology in Population B. Calculated FoldX energies were lowest for three candidate fragments of medin, with LQVDLGSSK being the most energetically favorable. However, the lower resolution of the extra density resulted in weaker correlation between energetic and map-fit metrics in our modeling approach (Supplementary Fig. 8), precluding us from assigning this density to a fragment of medin.

So, despite the identification of adjacent, low-resolution density in Population B fibrils, our modeling efforts were inconclusive in definitively assigning this density to a specific medin fragment. The limited local resolution, combined with the apparent structural heterogeneity, likely reflects flexibility or multiple conformations of the interacting species. Nonetheless, the consistent presence of this peripheral density across the reconstructed fibrils implies a role for unresolved, possibly transient heterotypic interactions in stabilizing this polymorph.

### Population B Fibrils Are Energetically Closer to Ex Vivo Aβ40 and Mutant Aβ42 Fibrils

To explore the structural context of Population B fibrils, we compared them to 26 previously reported Aβ fibril structures spanning Aβ40 and Aβ42 isoforms, derived from in vitro assembly, seeded reactions, and ex vivo extracts from patient or mouse brain tissue. For each structure, we computed residue-wise FoldX energy profiles to quantify stabilizing interactions and visualized their similarities using principal component analysis (PCA) followed by t-distributed stochastic neighbor embedding (t-SNE), a non-linear dimensionality reduction technique that preserves local structural relationships in two-dimensional space (Van der Maaten and Hinton 2008). This analysis provides an objective, energetically informed comparison of fibril architectures based on their underlying stabilizing interaction patterns, offering a structure-derived map that reflects how individual residues contribute to the overall fold stability.

In the t-SNE map (Fig. 4A), Aβ42 fibrils form a tight central cluster, suggesting that they represent a continuum of structural variants with similar stabilizing features. In contrast, Aβ40 fibrils are more widely dispersed, consistent with greater conformational diversity. Importantly, in vitro-derived Aβ40 fibrils tend to cluster on one side of the map, while ex vivo fibrils extracted from CAA brain cluster on the opposite side, suggesting they are stabilized through distinct structural mechanisms.

**Figure 4:**
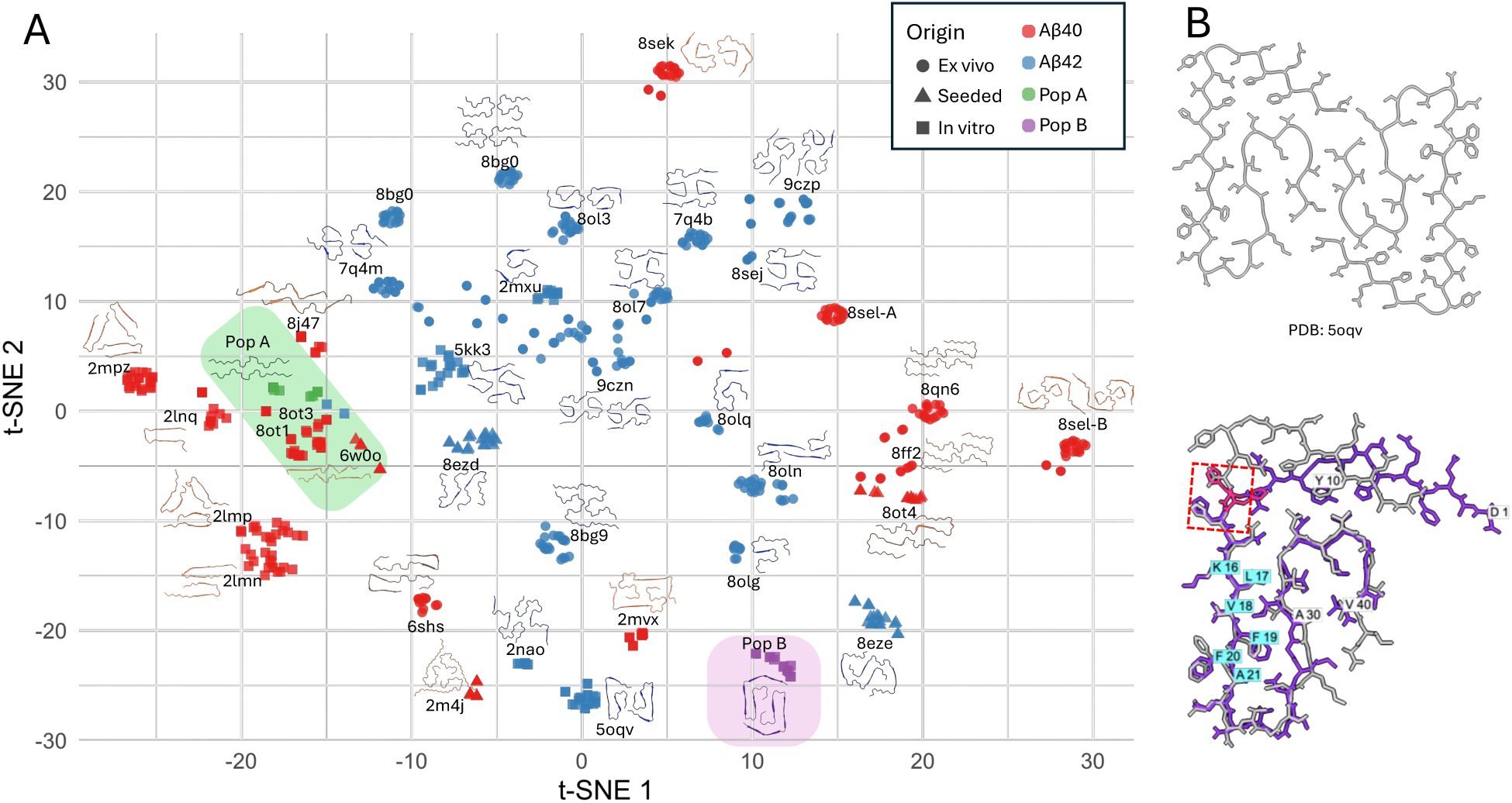
Population B structurally converges with Aβ42 fibrils. **(A)** t-SNE projection of Aβ fibril polymorphs based on per-residue FoldX energy profiles. Each point represents a published Aβ40 or Aβ42 fibril structure or a structure from this study (Pop A and Pop B). Colors indicate isoform and origin (e.g., in vitro, ex vivo, seeded). Population A clusters tightly with canonical in vitro Aβ40 structures. In contrast, Population B groups with a subset of Aβ42 fibrils, including 8olg (from APP23 mouse brain) and 5oqv (seeded Aβ42 fibrils), indicating conformational similarity. These fibrils share features such as a structured N-terminus, buried hydrophobic C-terminal loop, and polar, non-planar subunit geometry. Despite being composed of the Aβ40 sequence, the Population B morphology mimics key structural characteristics of Aβ42 fibrils and supports the view that heterotypic interaction with medin can shift Aβ40 folding toward an Aβ42-like energetic and architectural state. **(B)** Structural alignment of the F-subunit (violet) with the L-S subunit (grey) of in vitro produced Aβ42 (PDB ID: 5oqv). The two subunits are perfectly aligned from the C-terminus up to His14; deviation begins at His13 (highlighted by a red box).

A key discriminating factor is the N-terminal region (D1-Q15). Most in vitro Aβ40 fibrils lack ordered N-terminal segments and do not involve these regions in fibril stability. In contrast, ex vivo-derived Aβ40 fibrils from CAA cases (pdb : 8qn6, 6shs, 8qn7) (Kollmer, Close et al. 2019, Yang, Murzin et al. 2023) feature a structured N-terminus that contributes to fibril stability. These observations suggest that the environment in which Aβ aggregates -whether in vitro, in vivo, or in the presence of cofactors - plays a critical role in defining its final conformation.

Strikingly, Population B does not cluster with canonical in vitro Aβ40 fibrils but rather maps to a region closer to Aβ42 grown by seeding with AD brain extract (pdb : 8eze) (Lee, Yau et al. 2023) and adjacent to mutant Aβ42 fibrils extracted from APP transgenic mouse models (e.g., tg-SwDI , pdb : 8olg) (Zielinski, Peralta Reyes et al. 2023). This positioning is consistent with its structural features: Population B exhibits a well-resolved N-terminal region that engages in stabilizing interactions, a feature not typically observed in in vitro Aβ40 structures. Moreover, the presence of peripheral cryo-EM density near the N-terminus - likely corresponding to a medin-derived segment - suggests that heterotypic co-aggregation with medin may directly stabilize this region.

The F-shaped fold of Population B resembles the L-S topology seen in Aβ42 fibrils produced in vitro (pdb : 5oqv; Fig. 4B) (Gremer, Schölzel et al. 2017), especially in the arrangement of the non-planar subunits and the hydrophobic C-terminal loop (Fig. 4B). However, their N-terminal regions diverge significantly. In the L-S topology, residues D1–H13 are folded inward and stabilized by salt bridges involving H6, H13, and E11 (Gremer, Schölzel et al. 2017). In Population B, this region remains open and solvent-exposed, and H13 is no longer available for salt bridge formation. Instead, K28 forms hydrogen bonds with the backbone of V40, and additional polar interactions involve D1, D23, and E22 in a distinctive zipper-like cluster (Fig. 3D). These differences highlight how small variations in charge interactions at the N-terminus can dramatically influence overall fibril morphology.

Taken together, our comparative analysis shows that Population B shares key stabilizing features with ex vivo Aβ40 and Aβ42 mutant fibrils, despite being formed from wild-type Aβ40 in vitro. Its distinctive N-terminal ordering and proximity to putative medin interaction sites suggest that heterotypic co-aggregation can direct Aβ40 into polymorphic states otherwise inaccessible under homotypic in vitro conditions.

### Population C: First atomic structure of full-length medin reveals a polar steric zipper

In the third Aβ40-medin experiment, we observed a substantial enrichment of a distinct fibril morphology (Population C; ∼18% of particles). These fibrils exhibit a crossover distance of ∼40 nm and a diameter of ∼8 nm and were also detected at low abundance in the medin-only sample used for chromium side-shadowing (Supplementary Fig. 4D). Chromium shadowing indicates that Population C fibrils are left-handed (Fig. 1F).

Helical reconstruction in cryoSPARC yielded a 2.76 Å map (FSC 0.143; Supplementary Fig. 5C). The fibril comprises two protofilaments related by pseudo-21 screw symmetry, with a helical twist of −2.008° and rise of 4.8 Å. The protofilaments enclose a central cavity of ∼1.3 nm in diameter along the fibril axis (Fig. 5). Model building and side-chain density unambiguously identify Population C as full-length medin (50 residues) (Fig. 5B). No additional peptide-like density is observed, and we find no evidence for incorporation of Aβ in this structure.

**Figure 5 :**
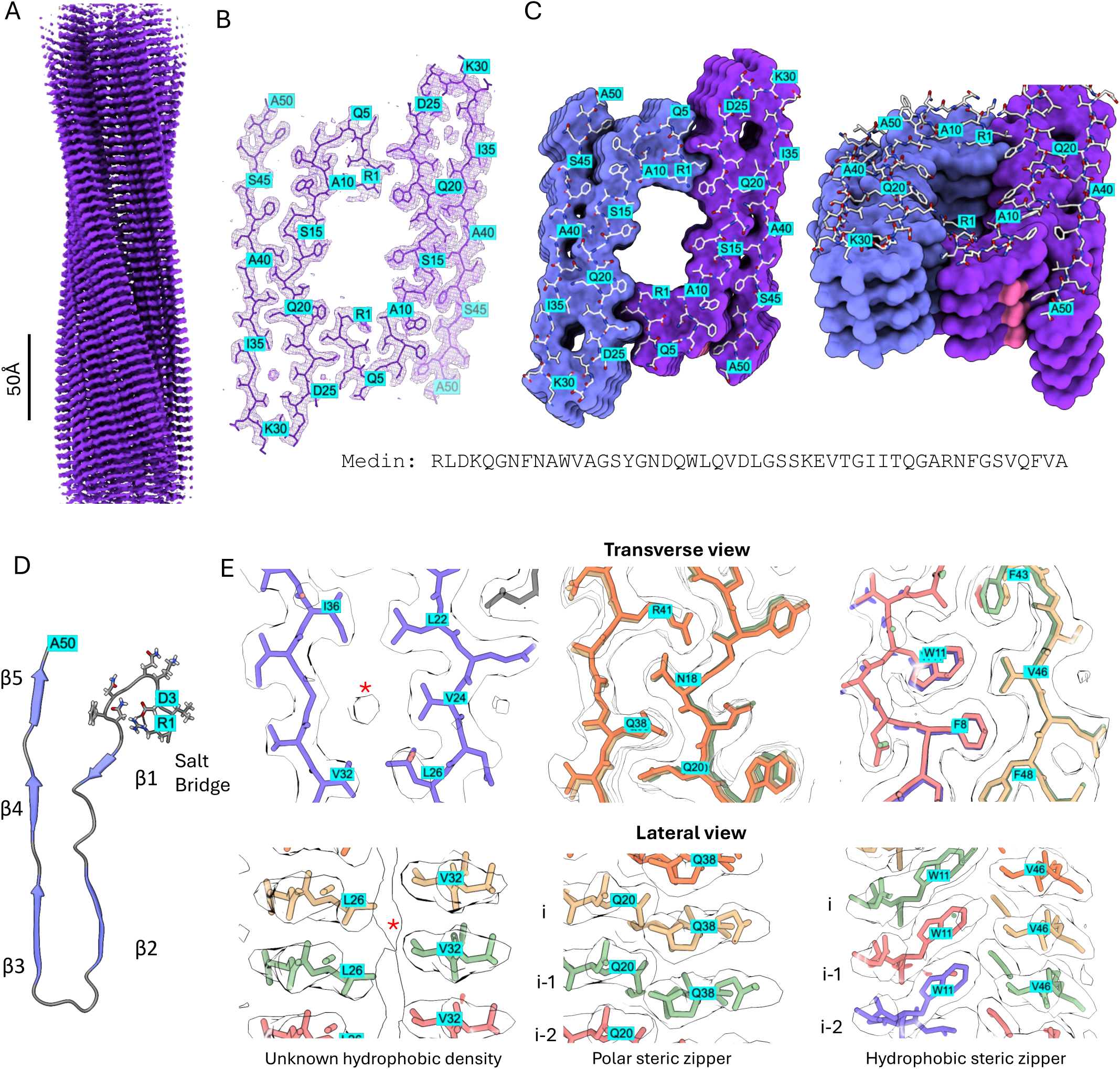
Population C reveals the first medin fibril structure with a polar steric zipper. **(A)** Cryo-EM density map of Population C fibrils reconstructed from the Aβ40–medin mixture (experiment 3). **(B)** Transverse section of the 3D cryo-EM density map fitted with two r-shaped subunits comprising medin R1–A50. The map has a mean of 0.0002058, s.d. 0.02169 (σ) and RMS 0.02169, and is displayed at a contour level of 5σ. **(C)** Surface representation of the atomic model of the Population C fibril, showing two protofibrils arranged in a staggered, polar configuration. **(D)** Cartoon representation of a single r-shaped medin subunit composed of five β-strands (β1–β5). Residues D3 and R1 (cyan) form an intramolecular salt bridge. Transverse and lateral section views highlighting stabilizing interactions within Population C fibrils. The chain-coloured atomic model is shown with the cryo-EM map outline. An additional density is observed within a hydrophobic pocket (4σ), surrounded by residues L22, V24, L26, V32, and I36. Residues N18, Q20, Q38, and R41 form a polar steric zipper between the N-terminal half of residue i and the C-terminal halves of residues i and i−1. Hydrophobic steric zipper interactions involving W11, F8, F43, V46 and F48 are formed between the N-terminal half of residue i and the C-terminal halves of residues i and i−2.

Each medin subunit adopts a distinctly non-planar, “r”-shaped conformation (Fig. 5D). Residues 1–26 form the buried fibril core and contribute to the inter-protofilament interface, whereas the C-terminal region is largely solvent-exposed. The subunit is organized into five β-strands: β1 (W11–A13), β2 (D19–D25), β3 (V32–T37), β4 (A40–F43), and β5 (V46–V49), with pronounced glycine-mediated kinks at G14 and G17. The N-terminus (R1–A10) forms a polar loop stabilized by a salt bridge between R1 and D3 (Fig. 5D). A hydrophobic loop connects β2 and β3, spanning residues V24 to I36. Interestingly, this region coincides with the β-hairpin structure proposed in the previously reported solid-state NMR model (Davies, Madine et al. 2015). This loop forms a hydrophobic pocket (∼8 Å in diameter) lined by V24, L26, and I35, within which an additional oval-shaped density is observed (Fig. 5E). Its identity remains unknown; however, the hydrophobic environment argues against water and is consistent with a small molecule bound with partial occupancy.

Protofilament stability is mediated by extensive steric-zipper interfaces. Notably, medin forms a polar steric zipper in which N18 and Q20 engage Q38 and R41 from neighboring subunits (Fig. 5E). In parallel, hydrophobic steric-zipper packing involves F9 and W11 from subunit i against F43, V46, and F48 from subunit i−2 (Figs. 5C, E). Protofilament association is further stabilized by a tightly interdigitated interface in which W21 and Q23 from one protofilament pack against L2 of the opposing protofilament (Fig. 5E).

## Discussion

Aβ fibrils are highly polymorphic, and increasing evidence links this diversity to disease context and biological activity. Yet most structural work has focused on homotypic assembly, leaving open how polymorphism is reshaped when multiple amyloidogenic species coexist. Here, using a defined Aβ40–medin mixture, we resolve atomic structures for fibrils derived from both components and show that mixed systems can produce coexisting “pure” fibrils alongside emergent polymorphs that are not observed under comparable homotypic conditions.

Our cryo-EM analysis identifies three fibril populations: Population A corresponds to a recurrent Aβ40 polymorph previously observed in vitro and also present in Aβ40-only samples, consistent with a common structural endpoint of Aβ40 assembly. In contrast, Population B is a previously unreported Aβ40 polymorph that emerges reproducibly in mixtures but not in Aβ40-only preparations. Population C is composed exclusively of medin and provides the first atomic structure of full-length medin fibrils. Together, these observations argue that mixed amyloid environments do not simply yield a single “hybrid” product; instead, they can expand the polymorphic ensemble, shift relative populations, and enable conformations that are otherwise disfavored.

A defining feature of Population B is its structured N-terminus, which is unusual for in vitro Aβ40 fibrils. In energetic space, Population B clusters away from canonical in vitro Aβ40 structures and closer to Aβ42 and ex vivo-like Aβ architectures characterized by ordered terminal regions. This supports the interpretation that the mixture reshapes the Aβ40 folding landscape toward more Aβ42-like structural solutions. The structured C-terminus in Population B reinforces this convergence: residues 28-40 form a compact hydrophobic loop and a stabilizing protofilament interface, features that resemble core-packing principles frequently observed in Aβ42 polymorphs despite the absence of residues 41-42. Together, these data show that Aβ40 can access Aβ42-like architectural states when assembled in a mixed amyloid environment.

What is the mechanistic basis of this remodeling? Biochemical and immunogold data indicate that Aβ and medin are associated within mixed assemblies, and Population B consistently exhibits peripheral density adjacent to the Aβ core. However, this density cannot be assigned at atomic resolution. Rather than viewing this solely as a technical limitation, it is consistent with a broader principle emerging from structural studies of amyloids: determinants that bias polymorph selection need not be rigidly ordered in the final structure. Heteromeric filaments such as patient-derived TDP-43/Annexin-11 demonstrate one end of the spectrum, where heterotypic interactions are fully ordered and structurally incorporated (Arseni, Nonaka et al. 2024). Our data are consistent with an intermediate regime in which heterotypic interactions act transiently and/or heterogeneously yet still leave a reproducible structural imprint on the dominant Aβ fold. Thus, mixed amyloid systems may remodel polymorphism through transient templating interactions, partial incorporation, or a combination of both.

The medin structure provides an additional conceptual link to this framework. Population C reveals a non-planar “r”-shaped fold stabilized by both hydrophobic interfaces and an unusual polar steric zipper, highlighting that medin can form highly specific, non-canonical stabilizing interactions. Interestingly, comparison between the medin protofilament interface and the putative interaction site in Population B reveals a notable similarity. In the medin fibril, protofilament association involves residues L2 and K4 from one subunit interacting with Q23 from the opposing protofilament (Supplementary Fig. 8D). In Population B, although the resolution of the peripheral density does not permit definitive sequence assignment, the highest-ranking medin fragment in our modeling analysis localizes near Aβ residues V18 and K16, with Q23 of the modelled medin positioned at the center of the interface (Supplementary Fig. 8C). The recurrence of closely related residue clusters in both contexts is striking. While this observation does not constitute mechanistic proof, it suggests a potential structural compatibility between medin–Aβ contacts and the intrinsic medin protofilament interface. One speculative possibility is that docking of medin onto the open N-terminal region of Aβ in Population B could preconfigure medin in a geometry that subsequently favors its own self-association into the polar steric zipper observed in Population C. Although this remains hypothetical, the parallel use of similar residue clusters in both assemblies highlights how shared interaction motifs may underlie compatibility in mixed amyloid systems.

Finally, we emphasize two scope points. First, the relative abundance of fibril morphologies varies between experiments (Supplementary Fig. 4), reflecting the intrinsic heterogeneity of Aβ assembly and sensitivity to sample handling and grid conditions. Importantly, this variability does not alter the central observation that Population B emerges reproducibly in mixtures (Supplementary Fig. 4C). Second, while Population B exhibits striking structural differences, we did not assess its cytotoxicity, seeding efficiency, or in vivo relevance.

Determining whether mixture-induced polymorph remodeling translates into altered biological activity will require future work, ideally including structural characterization of native deposits from systems where medin and Aβ co-accumulate.

Overall, our findings establish that interactions between coexisting amyloids can remodel polymorphism in a mixed system, yielding emergent architectures while retaining pure fibril species. This provides a structural framework for understanding how heterotypic amyloid environments expand conformational space and may shape amyloid strain selection in ageing-associated pathology.

## Methodology

### ThT kinetics assay

Medin peptide was custom synthesized by Bachem and supplied as a lyophilized powder or synthesized in-house using the protocol described in De Vleeschouwer M. et al 2025

(De Vleeschouwer, Pradhan et al. 2025). To eliminate any pre-formed oligomers, the peptide was dissolved in 100% hexafluoroisopropanol (HFIP) in a 2 mL glass vial and incubated at room temperature for 10 minutes. The HFIP was then evaporated by applying a steady stream of nitrogen gas, leaving medin as a thin film on the vial walls. To prepare a stock solution of monomeric medin, this film was dissolved in Milli-Q (MQ) water. Aβ40 peptide was obtained from rPeptide (Catalog ID: A-1153) in HFIP-treated film form. The stock solutions of both medin and Aβ40 were filtered using Costar® Spin-X® Centrifuge Tube Filters (0.22 µm, Costar 8160) before being diluted to a final concentration of 20 µM in a buffer containing 50 mM Tris (pH 7.4), 100 mM NaCl, and 10 µM ThT. The prepared samples were transferred to μClear medium-binding half-area plates (Greiner, #675096) for fluorescence measurements. ThT fluorescence was monitored over 100 hours using a Fluostar fluorescence plate reader (BMG Labtech) at 30°C, with excitation and emission wavelengths set to 440 nm and 480 nm, respectively. Readings were recorded every 10 minutes, with 10 seconds of shaking prior to each measurement. Fluorescence data were analyzed and plotted as a function of time using GraphPad Prism.

### Immunoprecipitation of fibrils

Amyloid fibrils of Aβ, medin and medin + Aβ (each 20µM after 7 days at 30°C) were immunoprecipitated (IP) using standard procedures with Protein G magnetic beads (Dynabeads, Thermo Fisher Scientific). Samples from the ThT assay were first centrifuged at 21k rcf to separate the fibrils from any remaining monomers. Fibrils taken from the pellet were then incubated with Dynabeads (Thermo Fisher Scientific, Cat number: 10003D) conjugated to a medin (1H4) or Aβ (6E10) monoclonal antibody (10µg antibody/mg beads) for 1 hour at room temperature. Afterwards, the beads were washed four times with PBST (PBS + 0.05% Tween). .Bound fibrils were eluted from the beads by incubation in 100 mM glycine (pH 2) for 10 min at 16°C and immediately neutralized with 1 M Tris (pH 8.3). For downstream immunoblotting, eluates were denatured in loading buffer (10% glycerol, 2% SDS, 100 mM DTT, 0.1 M Tris-HCl, pH 8.6), heated to 95°C for 5 min, and sonicated (3x30 seconds, 4°C, Bioruptor Pico).

### Western blotting

Gel electrophoresis was performed as previously described (Wagner, Degenhardt et al. 2022). Briefly, proteins were separated on a Bis-Tris 4–12% gradient gel in MES SDS Running Buffer (NuPage, Invitrogen) and transferred to a nitrocellulose membrane in a semi-dry blotting system. Transfer was confirmed by Ponceau-S staining. For the detection of medin and Aβ, membranes were boiled in PBS for 5 minutes at 90°C. After blocking in 5% milk in PBST for 1 hour, membranes were incubated overnight at 4°C with the primary antibody in PBST. For sequential incubation, the primary antibodies anti-human medin 1H4 (2µg/ml), followed by anti-amyloid-β 6E10, were used. Membranes were probed with the respective secondary horseradish peroxidase (HRP)-labelled antibodies (1:20,000, Jackson Immuno Laboratories). Protein bands were detected using chemiluminescent peroxidase substrate (Super Signal West Pico Plus or Dura, Thermo Fisher Scientific).

### Analytical LC-MS

Liquid chromatography–mass spectrometry (LC-MS) analysis was carried out using a Shimadzu Prominence LC-30 system coupled with a Zorbax 300 Extend-C18 analytical column (2.1 × 100 mm, 3.5 µm, 300 Å) and a matching narrow-bore guard column (2.1 × 12.5 mm, 5 µm). Mass spectrometric detection was performed online with a Shimadzu LCMS-2020 single quadrupole detector operating in electrospray ionization (ESI) mode. The separation was conducted at 80°C with a flow rate of 0.35 mL/min, employing a binary solvent system consisting of 0.1% formic acid in water (solvent A) and 0.07% formic acid in acetonitrile (solvent B). Unless specified otherwise, detection was achieved using a photodiode array detector at a wavelength of 214 nm. LC-MS chromatograms for all samples were either plotted using GraphPad Prism or obtained directly from the proprietary software of the instrument.

### Immunogold labeling

A modified phosphate-buffered saline (PBS+), pH 8.2, containing 1% BSA, 500 µl/L Tween-20, 10 mM Na-EDTA, and 0.2 g/L NaN₃, was used in all subsequent procedures for antibody and secondary gold probe dilutions. Aliquots (4 µL) of the fibril samples were pipetted onto Formvar/carbon-coated 200 mesh copper TEM support grids (Agar Scientific, Essex, UK).

The grids were incubated for 2 minutes, after which excess liquid was removed with Whatman 1 filter paper. Grids were then blocked by placing on 20 µL drops of normal goat serum (1:10 in PBS+) for 15 minutes. Following blocking, grids were incubated with 20 µL drops of primary antibodies (6E10: Anti-Aβ from BioLegend^®^ catalogue ID 9320-02 and 1H4: Anti-medin, in-house generated ) at a final dilution of 1:100 for 2 hours at room temperature. Afterward, the grids were rinsed three times for 2 minutes each with PBS+ and then incubated with a gold particle-conjugated secondary antibody (1:50 dilution in PBS+) for 1 hour at room temperature. Grids were subsequently rinsed five times for 2 minutes each in PBS+, followed by five 2-minute rinses in distilled water. Finally, the samples were negatively stained with 1% uranyl acetate for 1 minute.

### Negative stain Transmission electron microscopy

After the ThT signal plateaued, fibrils from various reaction samples were collected and centrifuged at 21,000 rcf to pellet the fibrils. A 5 µL aliquot of the fibril suspension was applied to a glow-discharged EM grid (Formvar/Carbon on 400 Mesh Copper, AGAR SCI, AGS162-4) and left to adsorb for 3 minutes. Excess liquid was carefully blotted away using Whatman Grade 1 blotting paper. The grid was then sequentially washed with three 20 µL drops of Milli-Q water before being stained with 2% (w/v) uranyl acetate for 1 minute. After staining, excess stain was blotted off, and the grid was left to air dry. The prepared grids were examined using a JEM-1400 transmission electron microscope (JEOL, Japan) equipped with a LaB₆ filament and operated at an accelerating voltage of 80 kV. Images were acquired using an Olympus Quemesa camera.

### Chromium side shadowing

Fibril handedness was assessed by oblique chromium-shadowing TEM. Fibril suspensions (5 µL) were adsorbed onto glow-discharged EM grids (Formvar/Carbon on 400-mesh copper, AGAR SCI, AGS162-4). After a 3-minute incubation, excess sample was blotted with Whatman filter paper, and grids were washed three times with 10 µL of Milli-Q water before being air-dried for 30 minutes. A ∼2 nm chromium layer was then deposited at a 30° incident angle using a Leica ACE600 sputter coater in single-direction mode. Oblique metal deposition generated an asymmetric chromium ridge along each filament, which was subsequently visualized by TEM as described above. Handedness was determined from the orientation of this ridge relative to the fibril axis, enabling unambiguous discrimination between left- and right-handed twists.

### Cryo-EM Grid Preparation and Data Acquisition

In total, five datasets were collected (Supplementary Fig. 4): two from Aβ40-only samples (Experiments 1 and 3) and three from Aβ40–medin mixture samples (Experiments 1, 2, and 3). For Experiments 1 and 2, Cryo-EM grids were prepared using Quantifoil holey carbon grids (R1.2/1.3 Cu 300 mesh) that were glow-discharged for 60 s at 5 mA using an ELMO glow discharger (Cordouan Technologies). Sample aliquots (3 μL) were applied to the grids using a Gatan CP3 cryo plunger under controlled conditions (22 °C, 98% relative humidity), incubated for 1 min, and blotted from both sides for 3 s using Whatman No. 2 filter paper.

Grids were plunge-frozen in liquid ethane cooled by liquid nitrogen. For Experiment 3, R2/1 Quantifoil grids with a 2 nm carbon support layer were used. Grids were prepared using a Vitrobot (Thermo Fisher) with a 45 s wait time and 3 s blotting time, followed by plunge-freezing in liquid ethane cooled by liquid nitrogen.

Cryo-EM data for Experiments 1 and 2 were collected on a JEOL CryoARM300 microscope operated at 300 kV, equipped with a Gatan K3 Summit direct electron detector and an in-column Omega energy filter (20 eV slit width). Automated collection was performed using SerialEM v3.8.0 (Mastronarde 2003). Experiment 3 data were collected on a Titan Krios (Krios 1) operated at 300 kV, equipped with a Falcon 4i direct electron detector in counting mode, and automated data collection was performed using EPU v3.11.

Micrographs from Experiments 1 and 2 were recorded at a calibrated pixel size of 0.71 Å (CryoARM300), with exposure times of 2.796 s, dose-fractionated into 60 frames, resulting in cumulative doses of ∼60 e⁻/Å² (∼1 e⁻/Å² per frame). For Experiment 3, movies were recorded at a pixel size of 0.74 Å, nominal magnification of 165,000×, spot size 8, illuminating an area of 0.58 µm. Each movie was dose-fractionated into 31 frames over 2.67 s, resulting in a cumulative dose of ∼40.7 e⁻/Å² (∼1.31 e⁻/Å² per frame). Images were collected over a defocus range of −0.5 to −1.5 µm for CryoARM300 (Experiments 1 and 2) and −0.7 to −1.8 µm for Experiment 3.

### Image Preprocessing and Helical Reconstruction

All datasets were initially processed in CryoSPARC v4 to rapidly assess fibril composition using preprocessing tools, including Patch Motion for motion correction and Patch CTF for CTF estimation. Fibrils were traced using the CryoSPARC filament tracer tool with an 18 Å inter-box distance and extracted into particle boxes of ±590 Å. Multiple rounds of 2D classification were performed to remove carbon edge classes and poor-quality filament segments. To estimate the relative population distribution, the number of particles belonging to 2D classes corresponding to each fibril morphology was quantified after excluding obvious junk classes, as reported in Supplementary Fig. 4.

Helical reconstruction of Population A fibrils was performed in RELION 5. Raw movie frames were aligned and dose-weighted using MotionCor2 (Zheng, Palovcak et al. 2017), and CTF estimation was carried out using CTFFIND4 (Rohou and Grigorieff 2015).

Micrographs with poor CTF fits or excessive drift were discarded before particle picking. Fibrils were manually picked in RELION using the filament picker, excluding poorly defined or heavily bundled fibrils. Filament segments were initially extracted with a 288 Å box size and 14.4 Å inter-box distance (∼95% overlap) from 2× binned datasets. Multiple rounds of 2D classification were performed to remove noisy, aggregated, or discontinuous segments.

Coordinates from well-resolved classes were re-extracted using a 720 Å box size with 4× binning to capture the full crossover length. Additional rounds of 2D classification were performed to remove curved fibrils. A representative class displaying a complete crossover was selected for initial model generation using the relion_helix_inimodel2d tool. Subsequent 3D classification was carried out with helical reconstruction enabled, allowing optimization of helical rise and twist. Selected classes with well-resolved features were used for 3D auto-refinement, and helical symmetry was imposed based on the optimized parameters. Iterative CTF refinement (including anisotropic magnification correction) and Bayesian particle polishing were performed to enhance map resolution. Final reconstructions were sharpened and post-processed using DeepEMhancer (Sanchez-Garcia, Gomez-Blanco et al. 2021).

Fibrils of Populations B and C were processed in CryoSPARC v4.2. Multiple rounds of 2D classification were performed to obtain homogeneous particle classes. Helical refinement jobs in CryoSPARC were performed with the local optimization of twist and rise enabled, and the refinement was repeated multiple times to optimize helical parameters. 3D classification was used to remove poor particles, followed by additional local and global CTF refinement and helical refinement with the heterogeneous refinement option enabled.

All details regarding cryo-EM data collection parameters, helical reconstruction parameters, and map statistics are provided in Supplementary Table 1.

### Model Building and Refinement

Postprocessed maps were used for building the atomic model. For population B, the handedness was first inverted by flipping the map in the Z direction using ChimeraX (Goddard, Huang et al. 2018, Meng, Goddard et al. 2023). Initial atomic models for Aβ fibrils in Populations A and B were generated using ModelAngelo (Jamali, Käll et al. 2024), which was provided with the post-processed 3D density maps and a FASTA file containing the sequences of Aβ and medin. ModelAngelo successfully built Aβ backbone models for both populations. These preliminary models were manually inspected and adjusted in Coot (Emsley, Lohkamp et al. 2010), correcting for sidechain rotamers, loop conformations, and missing fragments. Real-space refinement was carried out in Phenix (Afonine, Poon et al. 2018), with secondary structure restraints applied where appropriate. For the lower-resolution density regions observed in Population A, peptide fragments were modeled as poly-alanine chains in Coot to mark uninterpretable stretches where sequence identity could not be confidently assigned. Attempts to model medin in these regions using ModelAngelo were unsuccessful, likely due to limited local resolution and conformational heterogeneity. All models were visualized and analyzed using UCSF ChimeraX and PyMOL. Final maps and models were validated using Phenix.real_space_refine, MolProbity (Davis, Leaver-Fay et al. 2007) to assess model geometry, rotamer outliers, and Ramachandran statistics. Resolution estimates were based on the gold-standard FSC (0.143 criterion), and local resolution maps were generated using RELION. Complete model validation metrics, FSC curves, and local resolution distributions are included in the Supplementary Information.

### Thermodynamic stability analysis of Aβ structures Using FoldX

To assess the thermodynamic stability and energetic contributions of individual residues to the amyloid fibrils, we employed the FoldX force field version 5.1 (1) for structural analysis of available cryo-EM models, as well as modelling the unknown densities. Amyloid fibril structures were retrieved from the Protein Data Bank (PDB) in .pdb format and were prepared for energy calculations using FoldX’s built-in RepairPDB function, which optimizes side chain rotamers and corrects common structural anomalies while maintaining the backbone conformation. Following structure repair, we applied FoldX’s Stability command to calculate the total folding free energy (ΔG) of each amyloid model. Calculations were performed at standard conditions (T = 298 K, ionic strength = 0.05 M, pH = 7.0) unless otherwise specified.

### Statistical analysis of FoldX energies

t-SNE plots were generated to compare the thermodynamic stability profiles of various Aβ structures. Before t-SNE, the numeric data were standardized (mean-centered and scaled to unit variance) and subjected to principal component analysis (PCA) using the prcomp function in R (v4.0.0). The top principal components were used as input to t-distributed stochastic neighbor embedding (t-SNE), implemented via the Rtsne package (v0.15) with a perplexity of 10 or 25, theta = 0.5, and dims = 2. A random seed of 123 was set to ensure reproducibility. T-SNE embeddings were visualized using ggplot2 (v3.4.0), with points colored by Aβ type and shaped by sample origin.

### Modeling unknown densities using FoldX

To model the unknown densities, we built a backbone trace into the density. We then used a sliding window approach over the sequence of the only two proteins in the sample; Aβ40 and medin and employed FoldX’s buildModel command to model the sidechains corresponding to each window, yielding a free energy estimate of each window as well as an atomic model.

Each model generated from FoldX was manually placed in the cryo-EM density and goodness of fit as determined using an inhouse pipeline. First, to refine the alignment between the electron density map and the atomic model, we applied a rigid transformation to maximize their fit. The quality of this fit was assessed using the Pearson correlation coefficient (L) between the theoretical volumetric occupancy of the atomic model and the electron density map. The optimization aimed to determine a 3D rotation matrix M (a 3×3 matrix uniquely defined by three angles) and a translation vector T (describing displacement in 3D space) that maximized L. Given the atomic coordinates X and the center C (calculated as the mean of X along each axis), the transformed coordinates were computed as: *X*′ = (*X* − *C*)*M* + *T* + *C*

We implemented this optimization using a gradient descent approach in PyTorch. The process began with initializing the atomic coordinates X and the electron density map E (400×400×400). The rotation matrix M was initially set as the identity matrix, while T was a zero vector, and C was computed as the mean of X. Using these parameters, the transformed coordinates X’ were calculated. Next, the theoretical atomic occupancy was generated using Pyuul (Orlando, Raimondi et al. 2022) in a differentiable manner, producing a 3D volume P with the same voxel size and resolution as E. To enhance computational efficiency, both E and P were cropped to the smallest bounding region that contained all voxels of P with values above 0.001. The voxels of P and E were then flattened into single-dimensional arrays, P_flat and E_flat, respectively.

The Pearson correlation coefficient L was computed using a differentiable function implemented in PyTorch. The gradients dL/dM and dL/dT were then determined using PyTorch’s automatic differentiation. The values of M and T were iteratively updated using the Adam optimizer with a learning rate of 0.0001, an epsilon value of 0.1, and AMSGrad enabled. This optimization process was repeated for 1,000 iterations, progressively refining M and T through gradient descent. The final output consisted of the optimized atomic coordinates and the maximized L value, providing an estimate of the consistency between the theoretical and experimental volumetric occupancy.

To create a map of the structural landscape of Aβ, we obtained all corresponding PDB structures (8qn7, 8qn6, 8sek, 8sel, 8ff2, 6shs, 2lmn2, lmp, 8j47, 2m4j, 8ot1, 8ot3, 8ot4, 6w0o, 2lnq, 2mpz, 2mvx, 2mxu, 2nao, 5kk3, 5oqv, 7q4b, 7q4m, 8bg0, 8bg9, 8ezd, 8eze, 8ol3, 8ol7, 8olg, 8oln, 8olq, 9czn, 9czp) as well as our own models and used the SequenceDetail command in FoldX to compute the contribution of each residue to the stability of the final structure for each chain in the structure. The matrix of all the energy profiles were used as input in a principal component analysis in R v3(prcomp command), and the tSNE function was used to generate a 2D project of this space that maximally conserves data structure (Rtsne function in the tsne library).

### Data Availability

The cryo-EM density maps for the fibril structures of Population A and Population B have been deposited in the Electron Microscopy Data Bank (EMDB) under accession codes EMD-54007 (Population A) and EMD-54006 (Population B). The corresponding atomic models have been deposited in the Protein Data Bank (PDB) under accession numbers 9RIW (Population A) and 9RIV (Population B). The density map and atomic model for Population C have been deposited under EMD-56480 and PDB 9U02, respectively. Mass spectrometry raw data have been uploaded to Zenodo and can be accessed at 10.5281/zenodo.18633076. Other source data are available from the corresponding author upon request.

## Acknowledgements

We would like to thank Marcus Fislage and Dirk Reiter at the Brussels Electron Cryo-Microscopy (BECM) facility for their invaluable support with cryo-EM data collection and analysis. We also acknowledge Frederik Delaere for managing the computational resources used in this study. Our gratitude extends to Georg Kislinger and Cornelia Niemann at the LMU Munich BMC Electron Microscopy (EM) facility for their assistance with TEM analysis. We further thank the Rega Institute, the eBIC at Diamond Light Source, and the University of Leeds Cryo-EM facility for their support with cryo-EM sample preparation and data collection.

BP is supported by a Junior Postdoctoral Fellowship (12AWB24N) from the Research Foundation – Flanders (FWO). This research was funded by grants from the FWO (ID G0C3522N) and the National Institutes of Health (NIH, Grant ID: R01AG079234), as well as Germany’s Excellence Strategy within the framework of the Munich Cluster for Systems Neurology (EXC 2145 SyNergy– ID 390857198). The content is solely the responsibility of the authors and does not necessarily represent the official views of the National Institutes of Health.

