## Supplementary figures and tables. for "Atomic structures of medin and Aβ fibrils reveal polymorphic remodeling in mixed amyloid systems"

[1] Switch Laboratory, VIB-KU Leuven Center for Neuroscience, 3000 Leuven, Belgium.
[2] Switch Laboratory, Department of Cellular and Molecular Medicine, KU Leuven, 3000 Leuven, Belgium.
[3] Biomedical Center (BMC), Biochemistry, Faculty of Medicine, LMU Munich, Munich, Germany.
[4] VIB-KU Leuven Center for Neuroscience, Biophysics Technology Unit, Herestraat 49, 3000 Leuven, Belgium
[5] Institute of Systems, Molecular & Integrative Biology, University of Liverpool, United Kingdom.
[6] Neuroimmunology and Neurodegenerative Diseases, German Center for Neurodegenerative Diseases (DZNE), Munich, Germany
[7] Department of Cellular Neurology, Hertie Institute for Clinical Brain Research, University of Tübingen, Tübingen, Germany
[8] Munich Cluster for Systems Neurology (SyNergy), Munich, Germany
[a] current address: National Centre for Cell Science, Pune, India
[b] current address: Center for Alzheimer's and Neurodegenerative Diseases, UT Southwestern Medical centre, Texas, USA
[c] current address: Laboratory of Pathogens and Host Immunity, University of Montpellier, CNRS and INSERM, Montpellier, France

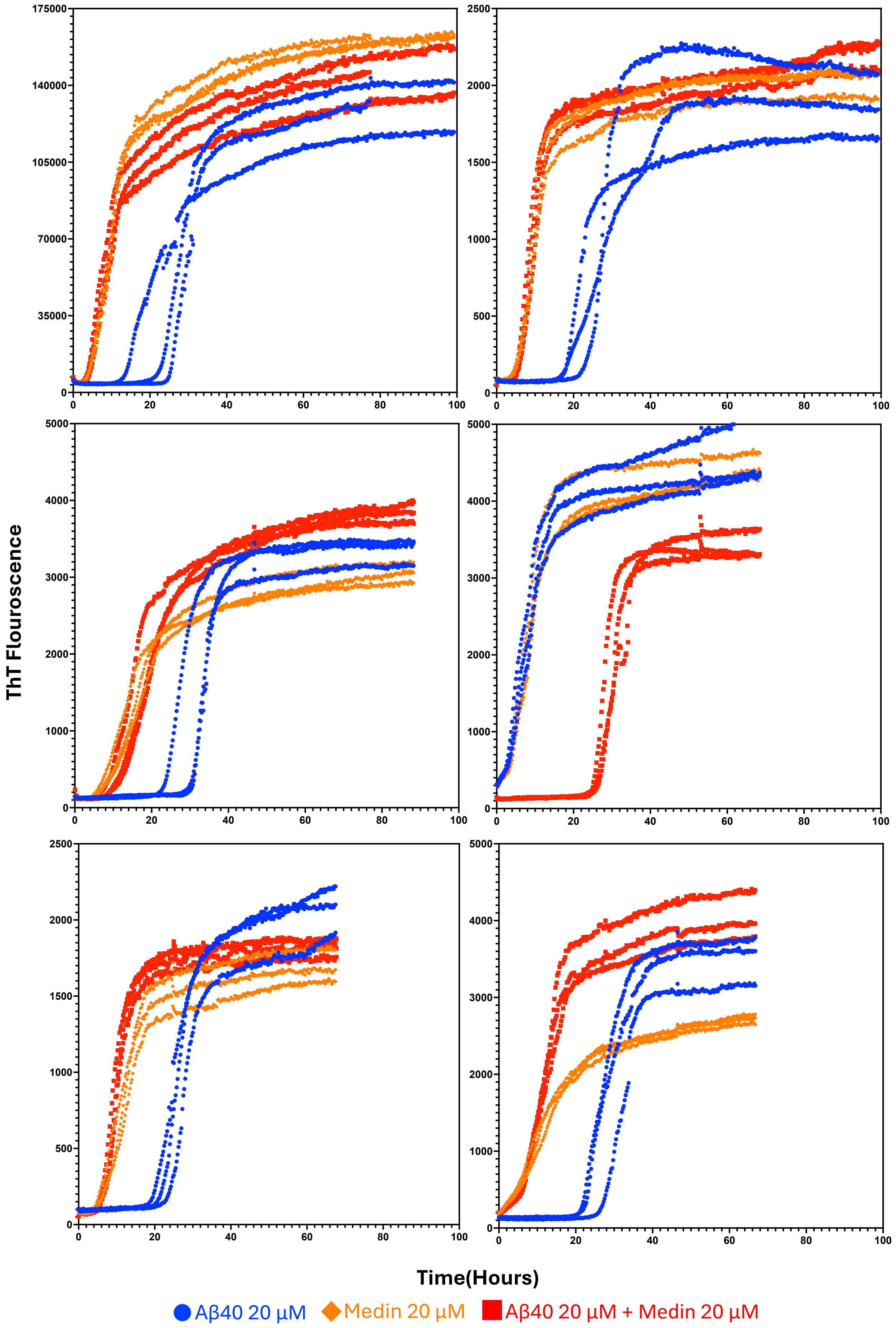

24 **Supplementary Figure 1: Independent ThT kinetics of Aβ40, medin, and their mixture.**

Six independent Thioflavin T (ThT) fluorescence kinetic assays of Aβ40 (20 μM), medin (20 μM), and an equimolar mixture were performed in 50 mM Tris, 100 mM NaCl, pH 7.4 at 30 °C. The kinetic curves were consistent across all independent experiments. Aβ40 exhibits a lag phase of ~23 h, whereas medin aggregates within ~3 h. The mixture shows an intermediate lag time (~6.5 h), indicating accelerated Aβ40 nucleation in the presence of medin. Each experiment included three technical replicates (n = 3). Raw fluorescence values are shown on the y-axis. Experiments were performed on two different plate readers with differing gain settings, accounting for the variation in maximum fluorescence intensity.

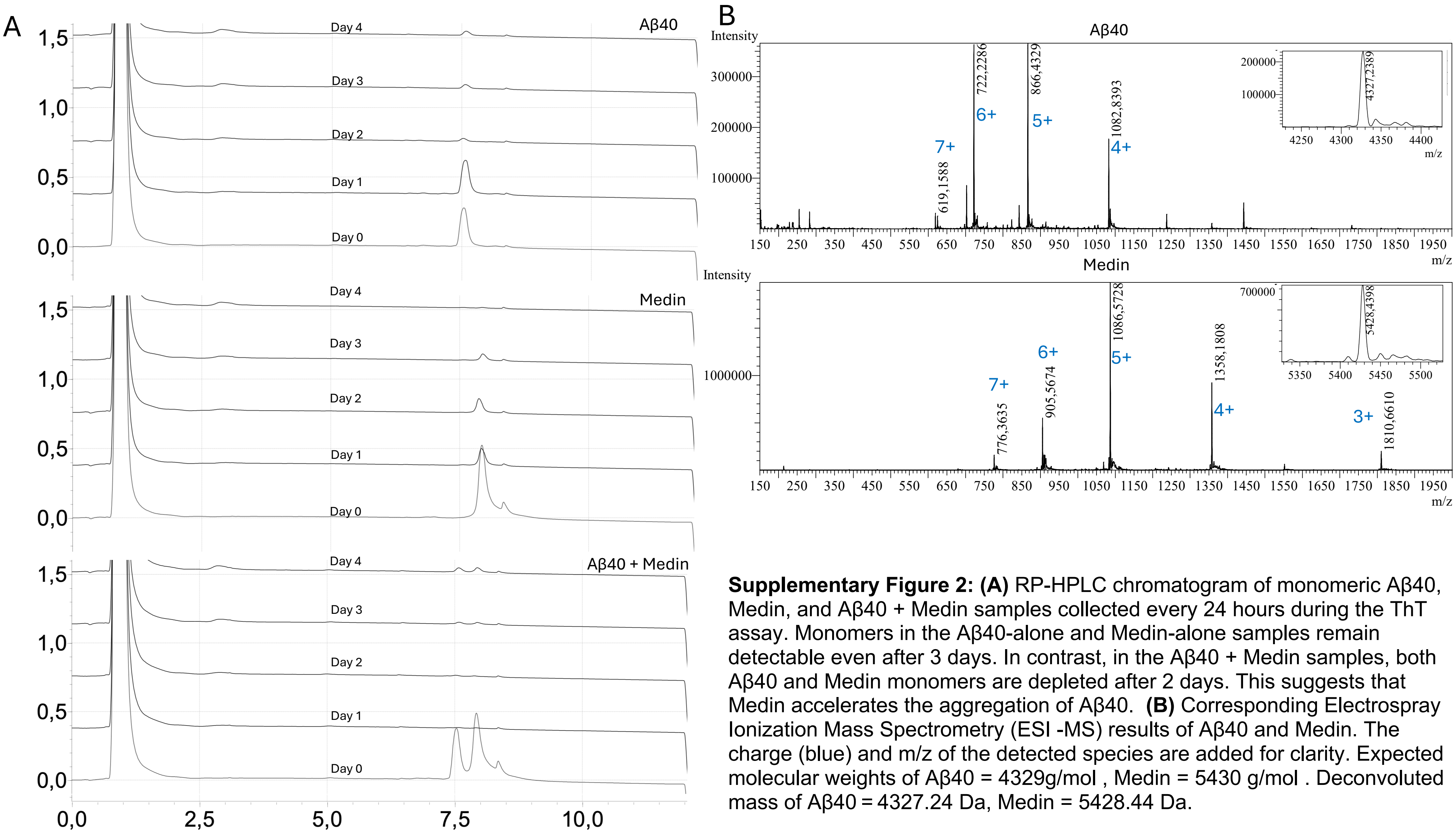

**Supplementary Figure 2: (A)** RP-HPLC chromatogram of monomeric Aβ40, Medin, and Aβ40 + Medin samples collected every 24 hours during the ThT assay. Monomers in the Aβ40-alone and Medin-alone samples remain detectable even after 3 days. In contrast, in the Aβ40 + Medin samples, both Aβ40 and Medin monomers are depleted after 2 days. This suggests that Medin accelerates the aggregation of Aβ40. **(B)** Corresponding Electrospray Ionization Mass Spectrometry (ESI -MS) results of Aβ40 and Medin. The charge (blue) and m/z of the detected species are added for clarity. Expected molecular weights of Aβ40 = 4329g/mol , Medin = 5430 g/mol . Deconvoluted mass of Aβ40 = 4327.24 Da, Medin = 5428.44 Da.

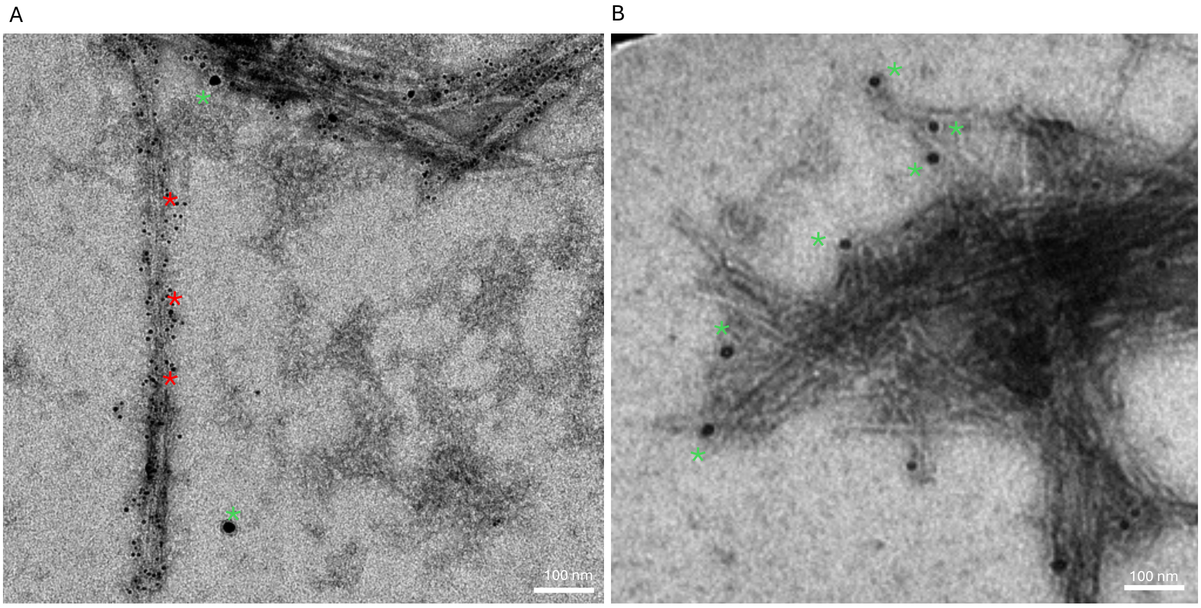

**Supplementary Figure 3:** Immunogold electron microscopy showing the presence of homotypic Aβ **(A)** and homotypic medin **(B)** fibrils in the Aβ40+medin sample (corresponding to Figure 1C). Dual immunogold labeling was performed using anti-Aβ (6E10; 6 nm gold, indicated by red asterisks) and anti-medin (1H4; 10 nm gold, indicated by green asterisks), confirming the coexistence of both fibril types in the mixture with heterotypic fibrils.

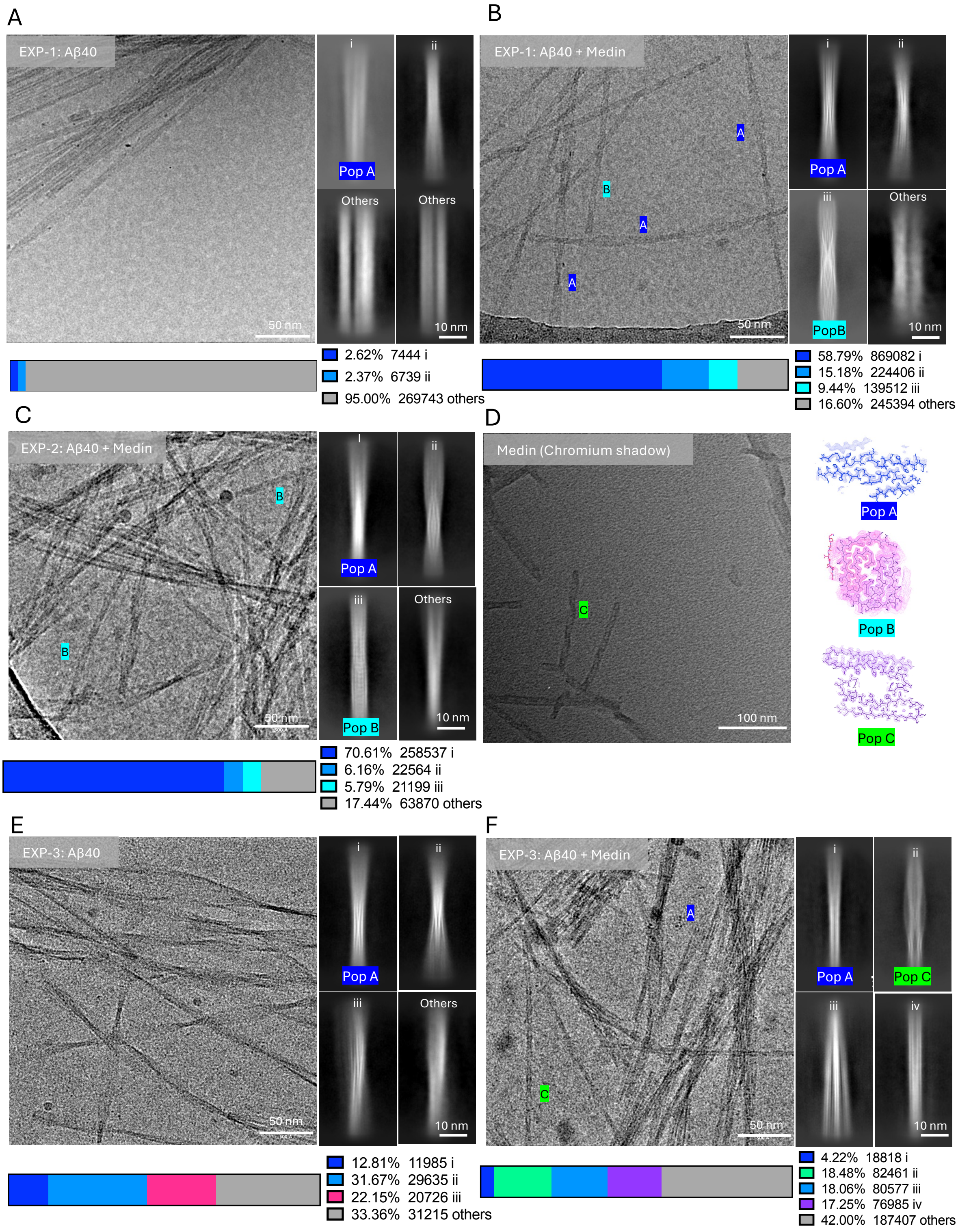

**Supplementary Figure 4: Morphology distribution of fibrils in Aβ40 alone and Aβ40–medin mixture.**

Five cryo-EM datasets (2 Aβ40, 3 Aβ40–medin mixture) were collected from three independent experiments. Shown are representative raw cryo-EM micrographs (left), 2D class averages of distinct fibril morphologies (right), and population distribution bars indicating both the percentage and absolute particle number for each morphology, labeled with Roman numerals I, II, III. Heterogeneous fibrils that do not belong to any defined class are designated as “others.” 2D classes were generated in CryoSPARC, with particles traced using the tracer tool at an 18 Å overlap and extracted with a box size of approximately ±590 Å. Population A is highlighted in blue, population B in cyan, and population C in green. Panels A and D show Aβ40-only samples, while panels B, C, and F show Aβ40–medin mixtures. Population A fibrils were observed in all datasets with varying proportions. Population B fibrils were detected only in experiments 1 and 2 (B, C) , whereas population C fibrils appeared only in experiment 3 (F). Grids for experiments 1 and 2 were prepared on Quantifoil R1.2/1.3 300 mesh holey grids, while experiment 3 used R2/1 grids with +2 nm carbon support (E,F). **(D)** Chromium-shadowed images of medin-only samples show the presence of population C fibrils even in the absence of Aβ40—thumbnails for each morphology on the right.

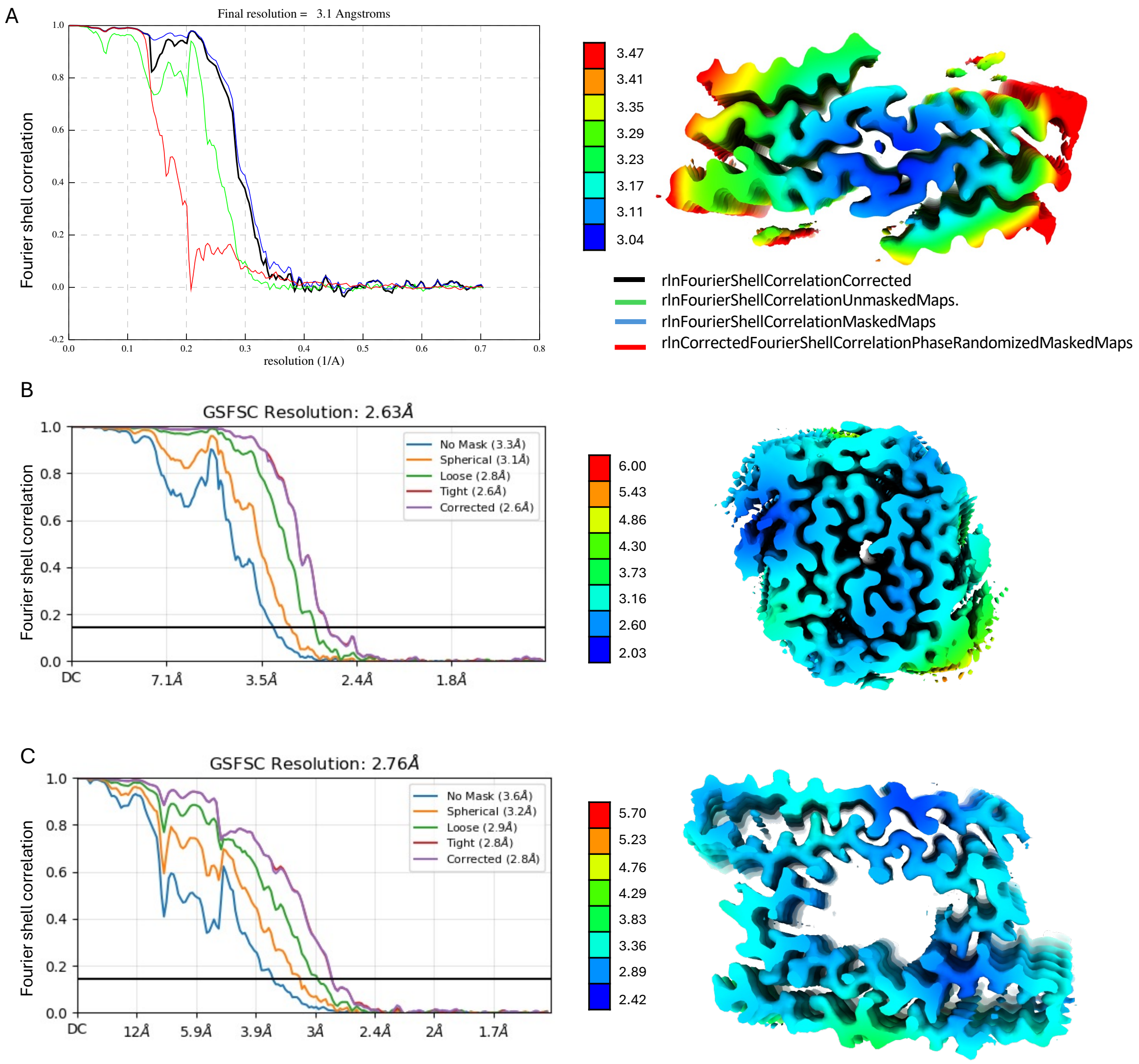

**Supplementary Figure 5 : Fourier Shell Correlation (FSC) curves and local resolution maps of cryo-EM helical reconstructions for fibril populations A, B, and C.** Population A was processed using RELION, while populations B and C were processed using CryoSPARC.

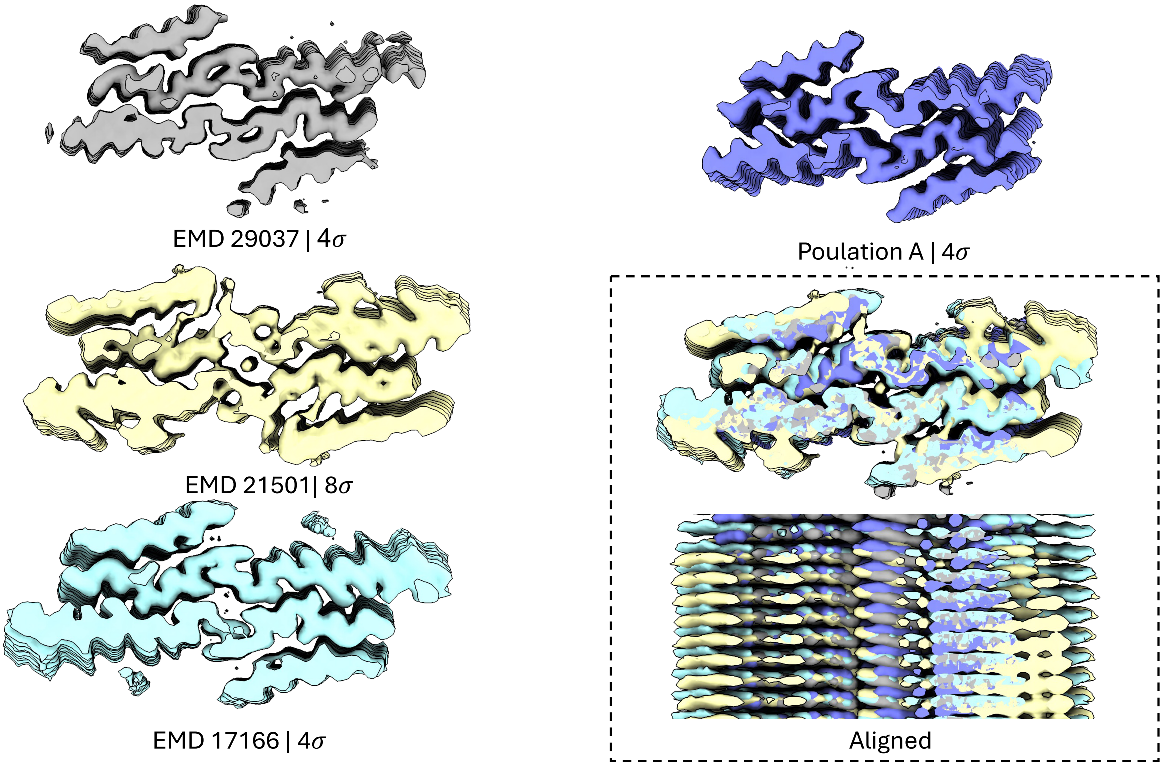

**Supplementary Figure 6 : Comparison of cryo-EM map of Population A with of A$\beta_{40}$ fibrils reported in the literature.** Fibrils were grown either by seeding with patient-derived samples (EMD-29037, grey; EMD-21501, yellow)(Ghosh, Thurber et al. 2021, Fu, Crooks et al. 2024) or unseeded (EMD-17166, cyan)(Pfeiffer, Ugrina et al. 2024), and compared with population A from this study. All structures adopt a four-layered extended conformation, with the inner, higher-resolution layers successfully modeled, while approximately ±15 residues from the N-terminus are missing. The two outer β-sheet layers are of lower resolution and could not be reliably modeled. All maps were resampled to the same scale and pixel size and visualized at 4σ, except for EMD-21501, which was visualized at 8σ for proper comparison

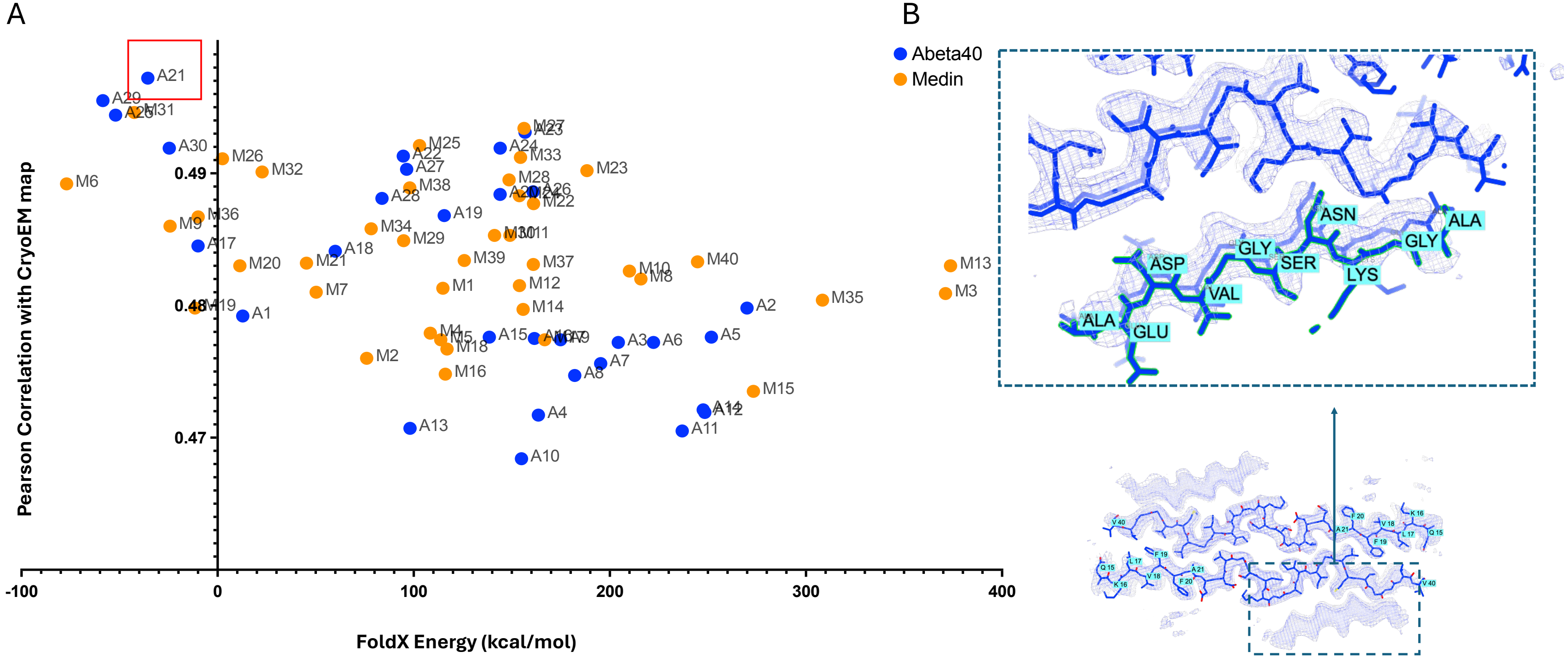

Aβ40:DAEFRHDSGYEVHHQKLVFFAEDVGSNKGAIIGLMVGGVV

**Supplementary Figure 7: The unknown density in Population A fibrils is likely a fragment of Aβ**. Scatter plots showing the Pearson correlation coefficient between the fit of FoldX- generated models (comprising fragments from Aβ40 and medin) and the unknown densities, plotted against stack energy (total folding free energy, ΔG) under standard conditions (T = 298 K, ionic strength = 0.05 M, pH = 7.0). Candidate fragments from Aβ40 are shown in blue, and those from medin in orange. The top-scoring candidate A21 = AEDVGSNKGA (Pearson coefficient = 0.4972, FoldX stack energy = –35.61 kcal/mol), is highlighted with a red box. The inset shows corresponding structural models fitted into the cryo-EM density, with the predicted sequences of the unknown densities highlighted in cyan.

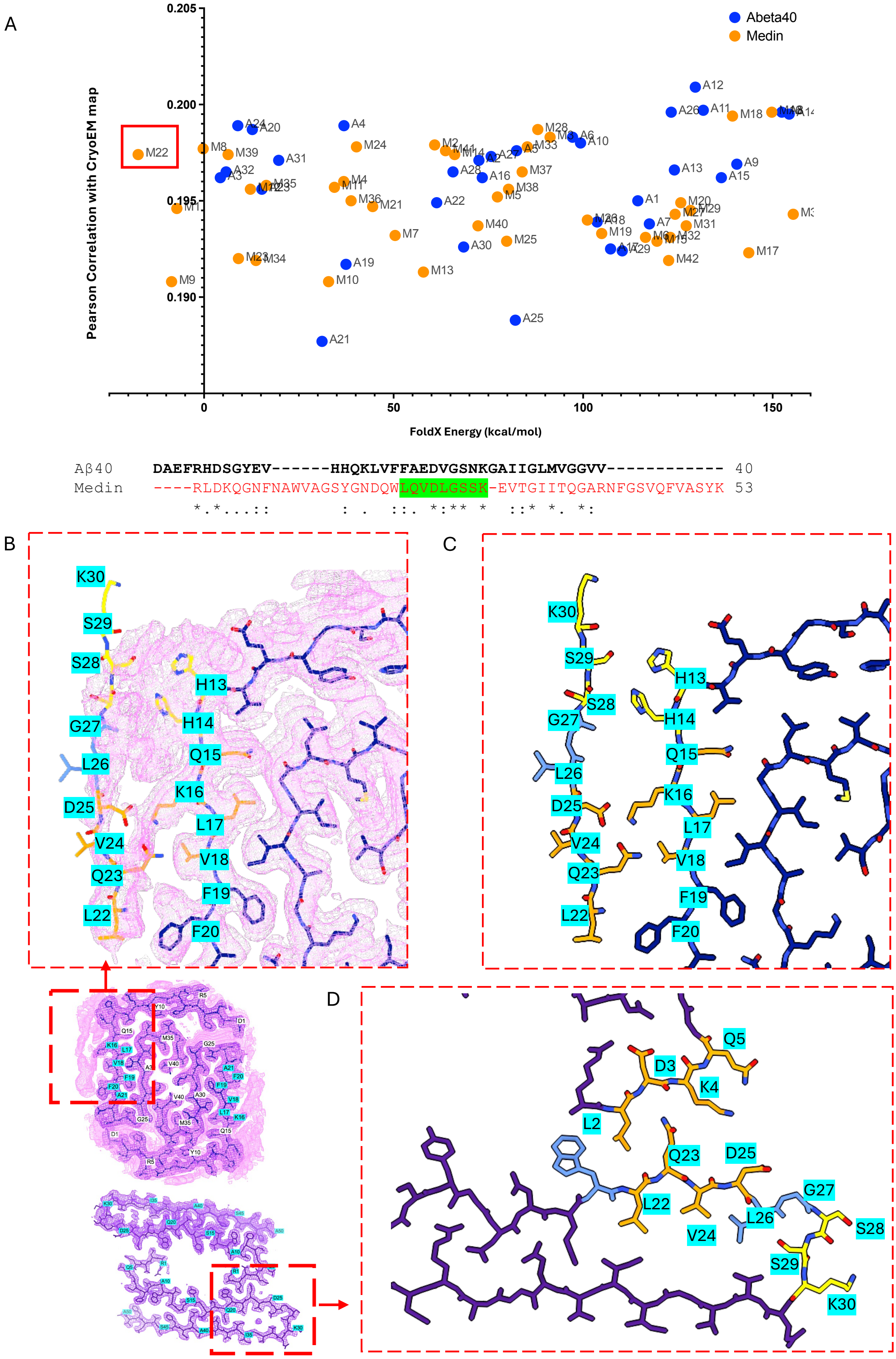
A
Abeta40
Medin
Pearson Correlation with CryoEM map
FoldX Energy (kcal/mol)
0.205
0.200
0.195
0.190
0
50
100
150
M22
Aβ40 DAEFRHDSGYEV------HHQKLVFFAEDVGSNKGAIIGLMVGGVV------------ 40
Medin ----RLDKQGNFNAWVAGSYGNDQWLQVDLGSSK-EVTGIITQGARNFGSVQFVASYK 53
B
C
D
K30
S29
S28
G27
L26
D25
V24
Q23
L22
H13
H14
Q15
K16
L17
V18
F19
F20
Q5
D3
K4
L2

**Supplementary Figure 8: The unknown density in Population B fibrils corresponds to a medin fragment that mimics the inter-protofibril steric zipper**. **(A)** Scatter plots showing the Pearson correlation coefficient between the fit of FoldX-generated models (comprising fragments from Aβ40 and medin) to the unknown cryo-EM density, plotted against stack energy (total folding free energy, ΔG) under standard conditions (T = 298 K, ionic strength = 0.05 M, pH = 7.0). Candidate fragments from Aβ40 are shown in blue and those from medin in orange. The top-scoring candidate for Population B, M22 (LQVDLGSSK; Pearson coefficient = 0.1974; FoldX stack energy = –17.40 kcal/mol), is highlighted with a red box. Sequence alignment of medin with Aβ40 is shown, with M22 highlighted in green, demonstrating high sequence similarity to the corresponding region of Aβ40. **(B)** FoldX-predicted structural model of the M22–Aβ40 complex fitted into the cryo-EM map of Population B.**(C)** FoldX-predicted model of the M22–Aβ40 complex highlighting key interface residues. Q23 of medin is proposed to form a steric zipper interaction with K16 and V18 of Aβ40 (shown in orange). A corresponding cryo-EM density consistent with this interaction is visible in panel B. **(D)** Inter-protofibril interface of the full-length medin structure, highlighting Q23 from one subunit forming a steric zipper with L4 and K4 of the opposing subunit from the adjacent protofibril. Panels B and C illustrate the structural similarity between the M22–Aβ40 interface and the inter-protofibril interface observed in Population C (full-length medin) fibrils.

**Supplementary Table 1**

### Cryo-EM data collection, refinement and validation statistics

| | Population A<br>EMD-54007<br>PDB 9RIW | Population B<br>EMDB-54006<br>PDB 9RIV | Population C<br>EMDB-56480<br>PDB 9U02 |
|---|---|---|---|
| **Data collection and processing** | | | |
| Magnification | 60000 | 60000 | 165000 |
| Voltage (kV) | 300 | 300 | 300 |
| Electron exposure (e–/Å$^2$) | 60 | 60 | 40.71 |
| Defocus range (μm) | 0.5-1.5 | 0.5-1.5 | 0.7-1.8 |
| Pixel size (Å) | 0.71 | 1.39 | 0.74 |
| Symmetry imposed | HELICAL, twist= -0.91, rise= 4.83 Å, axial sym = C1 | HELICAL, twist = 2.85, rise = 4.77 Å, axial sym = C1 | HELICAL, twist = -2.008, rise = 4.804 Å, axial sym = C1 |
| Initial particle images (no.) | 68372 | 160508 | |
| Final particle images (no.) | 34945 | 114792 | 47541 |
| Map resolution (Å)<br>FSC threshold | 3.10<br>0.143 | 2.58<br>0.143 | 2.76 |
| Map resolution range (Å) | 3.0 - 3.5 | 2.2-6 | 2.4-5.7 |
| **Refinement** | | | |
| Initial model used (PDB code) | NONE | NONE | NONE |
| Model resolution (Å)<br>FSC threshold | 2.1<br>0.143 | 1.9<br>0.143 | 2.3<br>0.143 |
| Model resolution range (Å) | | | |
| Map sharpening *B* factor (Å$^2$) | -63.3423 | -31.3 | -31.293. |
| Model composition<br>Non-hydrogen atoms<br>Protein residues<br>Ligands | <br>1480<br>208<br>0 | <br>3050<br>400<br>0 | <br>3840<br>500<br>0 |

| | | | |
|---|---|---|---|
| *B* factors (Å$^2$)<br>Protein<br>Ligand | Min-Max/Mean<br>55.35-128.49/77.09<br>NA | Min-Max/Mean<br>46.20-93.65/61.65<br>NA | Min-Max/Mean<br>31.59-82.23/49.04<br>NA |
| R.m.s. deviations<br>Bond lengths (Å)<br>Bond angles (°) | <br>0.006<br>0.772 | <br>0.002<br>0.346 | <br>0.003<br>0.467 |
| Validation<br>MolProbity score<br>Clashscore<br>Poor rotamers (%) | <br>1.25<br>0<br>0 | <br>0.89<br>1.51<br>0 | <br>1.49<br>1.73<br>0 |
| Ramachandran plot<br>Favored (%)<br>Allowed (%)<br>Disallowed (%) | <br>79.17<br>20.83<br>0 | <br>100<br>0<br>0 | <br>89.58<br>10.42<br>0 |
